# Persistent interferon signaling and clonal expansion mark early events in DNA methylation damage-induced liver cancer

**DOI:** 10.1101/2025.10.01.679856

**Authors:** Lee J. Pribyl, Jennifer E. Kay, Joshua J. Corrigan, Lindsay B. Volk, Monét Norales, Norah A. Owiti, Evan A. Kowal, Ishwar N. Kohale, Ilana S. Nazari, Matilda R. Swanson, Aimee C. Moise, Duanduan Ma, Stuart S. Levine, Emily Michelsen, Robert G. Croy, Timothy Ragan, Dorothea K. Torous, Svetlana L. Avlasevich, Stephen D. Dertinger, Sebastian E. Carrasco, Leona D. Samson, John M. Essigmann, Forest M. White, Bevin P. Engelward

## Abstract

N-Nitrosodimethylamine (NDMA), a probable human carcinogen, induces toxic and mutagenic *O*^6^-methylguanine lesions that are repaired by the *O*^6^-methylguanine methyltransferase (MGMT). To elucidate mechanisms of NDMA-induced liver cancer progression, we performed longitudinal analyses of phenomic, transcriptomic, and phosphoproteomic changes in wild-type and MGMT-deficient mice, observing amplified responses in the deficient genotype. Early molecular rewiring indicative of a DNA damage response was detected by phosphoproteomic and transcriptomic profiling within days post-exposure. Transcriptomic analyses identified a persistent and robust interferon response as the dominant activated pathway. This chronic interferon signaling, which remained unresolved, correlated with extensive clonal expansion, an early hallmark of oncogenesis. Spatial transcriptomics further revealed pathway alterations favoring tumorigenesis within clonally expanded cells. These findings delineate the cascade of molecular events triggered by acute early-life NDMA exposure, culminating in cancer development months later. Our study unveils potential predictive biomarkers and strategies for disease mitigation.

## Introduction

Alkylating agents are among the most abundant and most hazardous DNA damaging agents^1,2^. N-Nitrosodimethylamine (NDMA), classified as a Group 2A probable human carcinogen^1,3^, is a methylating agent responsible for the recall of over a dozen contaminated drugs^4–6^. Furthermore, NDMA contamination from chemical waste leaching at the Olin Chemical Superfund site has been associated with elevated rates of childhood cancer^7–9^. While it is well-established that NDMA induces cancer in animal models^6,10^, the nature and temporal dynamics of systems-level responses that modulate disease progression remain poorly defined. Moreover, methylating agents are used as chemotherapeutics^11^, making this study relevant to both cancer causation and treatment.

NDMA is primarily metabolized in the liver by cytochrome P4502E1 (CYP2E1), creating a methyldiazonium ion that results in DNA adducts. One of the most biologically impactful adducts is *O*^6^-methylguanine (*O*^6^MeG), which is mutagenic, toxic, and carcinogenic^1,6,10,12^. *O*^6^MeG pairs readily with thymine, causing G to A transition mutations^13–15^. Toxicity arises when mismatch repair (MMR) repeatedly excises the newly synthesized strand opposite *O*^6^MeG, creating persistent single-strand gaps^1,14,16–18^. This process, known as futile cycling, can cause replication forks to collapse when they encounter these gaps in the next S phase, producing toxic double-strand breaks (DSBs) that can be repaired by homologous recombination (HR)^19^. While mostly accurate, HR can lead to large-scale sequence rearrangement mutations, including insertions, deletions, and translocations^20^. Cells mitigate *O*^6^MeG toxicity and mutagenicity through the *O*^6^-methylguanine methyltransferase (MGMT), a suicide protein that transfers the methyl group from *O*^6^MeG to an active site cysteine residue^21,22^.

In several contexts, DNA damage has been shown to stimulate interferon (IFN) signaling, thus linking structural DNA changes to immune responses^23^. For example, cGAS senses fragments of DNA in the cytosol, activating STING to initiate a Type I IFN response, which can contribute to Type II IFN (IFN-γ) responses mediated by JAK/STAT1 signaling^24^. While essential for defense against infections, chronic IFN activation promotes carcinogenesis through sustained inflammation and immune dysregulation^25^. Although DNA-damaging agents are known to induce the IFN response *in vitro*^24,26^, much less is known about methylation damage and its potential to drive IFN signaling *in vivo*, a critical gap in knowledge that is relevant to cancer susceptibility.

To elucidate the consequences of unrepaired *O*^6^MeG in the liver, we studied MGMT-deficient (*Mgmt*^−/−^) mice. Previously, we found point mutation frequency in *Mgmt*^−/−^ NDMA-treated mice were about 4-fold more abundant than in wild-type (WT)^16^. Interestingly, the mutational spectra in both strains were nearly identical, dominated by GC→AT mutations. In the present work, we quantified over 40 molecular, cellular, and physiological phenotypes, including DNA damage, protein levels, sequence rearrangements, clonal dynamics, and histopathology. By integrating phenomics with analysis of phosphoproteomics and transcriptomics, we generated a comprehensive understanding of disease progression. Furthermore, using the RaDR-GFP mouse model, a transgenic system that detects large-scale sequence rearrangements indicative of mutations, we discovered extensive clonal expansion. Analysis of spatial transcriptomics in this model revealed insights into gene expression changes early in tumor development. Taken together, these results provide mechanistic insights into the progression from DNA insult to malignancy, identifying potential biomarkers for disease risk and opening avenues for novel mitigation strategies.

## Results

### MGMT deficiency exacerbates NDMA-induced liver injury and tumorigenesis

We used a well-established approach to induce tumorigenesis wherein WT and *Mgmt*^−/−^ neonatal mice were administered 3.5 mg/kg and 7.0 mg/kg of NDMA at 8 and 15 days old, respectively, via intraperitoneal injection (**Fig. 1a**)^12,27^. Ten months post-exposure, *Mgmt*^−/−^ mice exhibited a 9-fold increase in liver tumors per mouse compared to WT, and pathology revealed the presence of hepatocellular carcinomas and adenomas (**Fig. 1b-d**). Male mice were twice as susceptible to tumorigenesis compared to females (**Extended Data Fig. 1a**), consistent with previous findings^10,12,28^. Severe weight loss occurred in *Mgmt*^−/−^ mice due to hepatobiliary injury secondary to tumor burden (**Extended Data Fig. 1b**). These results demonstrate that MGMT deficiency critically enhances susceptibility to NDMA-induced liver tumorigenesis and hepatocellular toxicity.

**Fig. 1:**
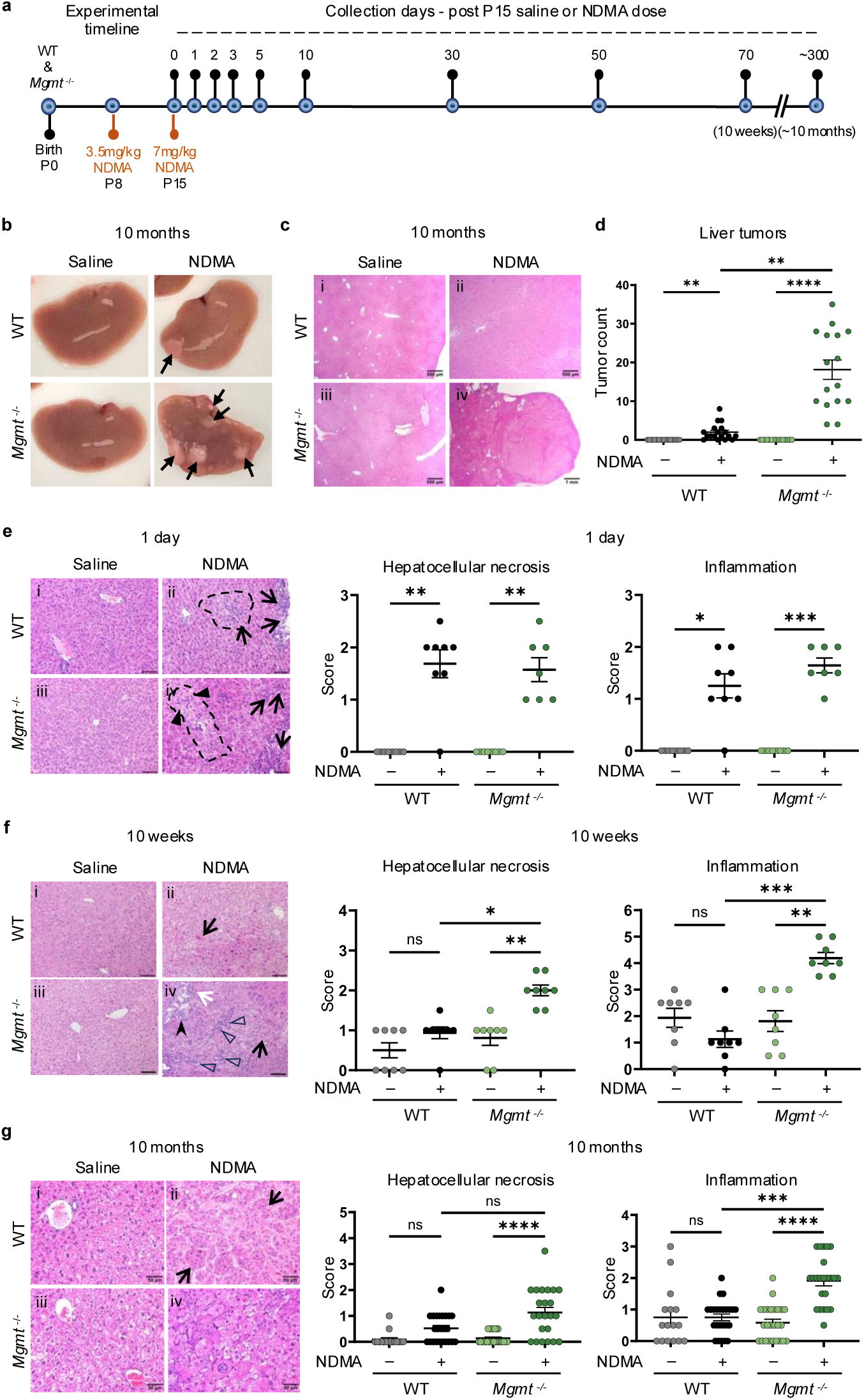
MGMT deficiency exacerbates NDMA-induced liver injury and tumorigenesis. **a**, Experimental timeline displaying two intraperitoneal injections of saline or NDMA at P8 (3.5 mg/kg) and P15 (7 mg/kg) in WT and *Mgmt*^−/−^ mice, with indicated collection timepoints. **b**, Representative gross images of livers from WT and *Mgmt*^−/−^ male mice 10 months post-treatment. Black arrows denote visible tumors. **c**, Representative H&E-stained liver sections at 10 months post-exposure (ii) show trabecular subtype hepatocellular carcinoma in NDMA-treated WT mice, and (iv) exhibit expansile neoplasms consistent with hepatocellular carcinoma (solid subtype) in *Mgmt*^−/−^ NDMA-treated mice (magnification 40x). Representative sections are depicted in photomicrographs in (**g**). **d**, Quantification of liver tumors at 10 months. *n* ≥ 15. **e**, Representative H&E sections (magnification 200x, scale bars = 100 µm) and histopathology scores for hepatocellular necrosis and inflammation at 1 day post-treatment. (ii) NDMA-treated WT mice showed periacinar hepatocellular degeneration (swelling; black dash line) with few macrophages and mild portal inflammation (arrows). (iv) *Mgmt*^−/−^ NDMA-treated mice exhibited focal/multifocal periacinar degeneration with or without single-cell necrosis (arrowheads) and mild portal/periportal inflammation (arrows). *n* ≥ 7. **f**, At 10 weeks post-NDMA treatment (ii) WT livers exhibited hepatocellular degeneration with increased cytoplasmic alterations, eosinophilia hypertrophy, and/or kariocytomegaly (black arrow). (iv) *Mgmt*^−/−^ mice exhibited multifocal hepatocellular degeneration (black arrow), with or without single-cell necrosis (white arrow), and mild to moderate lymphocytic and histiocytic inflammation in portal tracts (stealth black arrowhead). Biliary hyperplasia (black arrowheads) was common in portal tracts. Magnification 200x, scale bars = 100 μm. *n* ≥ 8. **g**, (ii) High-magnification H&E sections (400x) at 10 months (ii) show a hepatocellular carcinoma – trabecular subtype (arrows) in an NDMA-treated WT male mouse. (iv) An NDMA-treated *Mgmt^−/−^* mouse shows a hepatocellular carcinoma (solid subtype). Hepatocellular necrosis and inflammation scores show significant increases in *Mgmt^−/−^*NDMA treated mice compared to controls. *n* ≥ 16. Histopathology scoring was based on H&E stains across indicated panels (**e**–**f**). Data are presented as mean ± s.e.m. Statistical comparisons by Kruskal-Wallis and Dunn’s test (**d**–**f**). Statistical significance: *p < 0.05, **p < 0.01, ***p < 0.001, ****p < 0.0001. NS, not significant.

Given that aberrant tissue physiology can drive tumorigenesis^29,30^, we investigated the temporal progression of NDMA-induced liver pathology. While at 1 day post-exposure to the second dose, WT and *Mgmt*^−/−^ mice exhibited comparable levels of necrosis and inflammation (**Fig. 1e**), these phenotypes diverged by 10 weeks wherein *Mgmt*^−/−^ mice developed worsening pathology (**Fig. 1f**), alongside intensified biliary hyperplasia and fibrosis (**Extended Data Fig. 1c**). H&E analysis revealed a significant increase in karyomegaly in *Mgmt*^−/−^ livers (**Extended Data Fig. 1d**) and DAPI staining demonstrated enlarged nuclear size (**Extended Data Fig. 1e**), both of which remained elevated ten weeks post-exposure. In *Mgmt^−/−^* mice, the immune cell markers CD3ε and CD11c were elevated at 5 days and 10 weeks post-exposure (**Extended Data Fig. 1f**), reflecting sustained immune cell recruitment and activation. Consistent with liver injury, *Mgmt*^−/−^ mice showed increased spleen weight and decreased body weight at 10 weeks (**Extended Data Fig. 1g**). Unlike WT mice, at 10 months, *Mgmt*^−/−^ mice displayed advanced necrosis, inflammation, biliary hyperplasia, and foci of hepatocellular alterations, while karyomegaly remained elevated in both genotypes (**Fig. 1g & Extended Data Fig. 1h**). Thus, although WT and *Mgmt*^−/−^ mice showed similar initial pathology, the latter developed progressively more severe disease over time, culminating in malignant transformation.

### MGMT suppresses DNA damage retention, toxicity, and compensatory proliferation

To learn whether *Mgmt*^−/−^ mice suffer a higher burden of DNA adducts, we performed mass spectrometry. This revealed a ~3.5-fold higher accumulation of *O*^6^MeG lesions in *Mgmt*^−/−^ mice compared to WT at 5 days post-exposure (**Fig. 2a**). At 50 days, *O*^6^MeG levels were similar in WT and *Mgmt*^−/−^ mice, likely due to MGMT repair and dilution by cell proliferation, respectively. According to the futile cycling model, such lesions can yield DSBs via replication fork collapse. Consistent with this prediction, we observed phosphorylation of a key marker of DSBs and replication fork stress, gH2AX. Analysis of gH2AX immunofluorescence and protein levels revealed a significant increase, relative to saline controls, for both genotypes (**Fig. 2b-c & Extended Data Fig. 2a-c**). This result correlated with a significant rise in micronuclei (**Fig. 2d**), confirming DSB accumulation. Crucially, WT mice recovered rapidly, whereas *Mgmt*^−/−^ livers exhibited persistently elevated gH2AX levels (**Fig. 2b**) with punctate foci detectable up to 50 days post-exposure (**Fig. 2c**). These findings indicate continuous replication fork stress and DSBs, despite remediation of *O*^6^MeG adducts (**Fig. 2a**), suggesting a role for inflammation-induced DNA damage caused by reactive oxygen and nitrogen species.

**Fig. 2:**
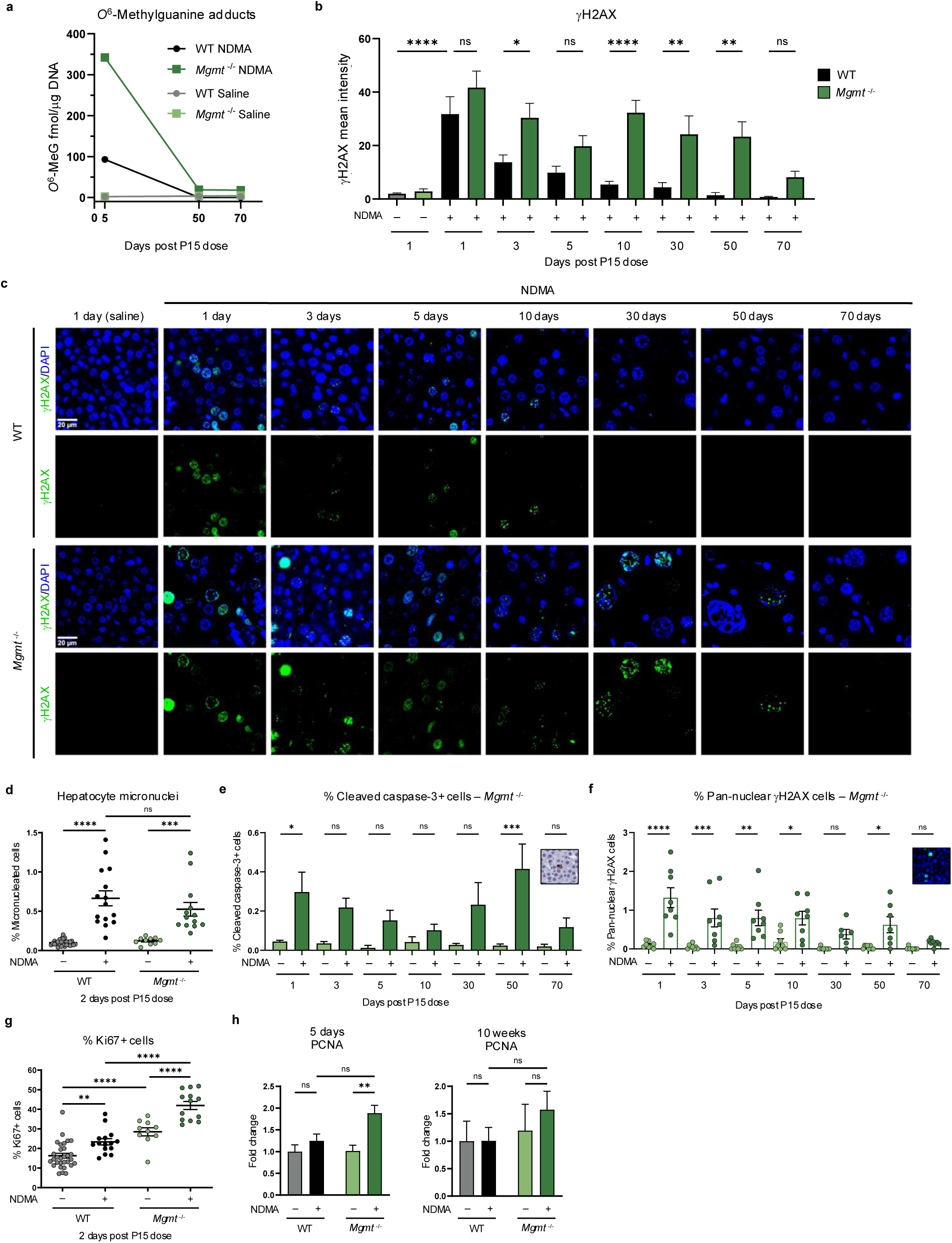
MGMT suppresses DNA damage retention, toxicity, and compensatory proliferation. **a**, Genomic DNA from liver tissue was analyzed for *O*^6^-methylguanine DNA adducts by triple quadrupole mass spectrometry. Adduct levels (fmol) were normalized to DNA (µg). Data points represent 1 male and 1 female sample combined. Saline controls received a single dose at P15. **b**, Quantification of the mean gH2AX fluorescence intensity per nucleus in NDMA-exposed WT (gray/black bars) to *Mgmt^−/−^* (green bars) livers. *n* ≥ 7. **c**, Representative immunofluorescent images of gH2AX (green) and DAPI (blue) stained liver sections across indicated post-treatment time points. Persistent DNA damage (gH2AX foci) and cell enlargement are observed in *Mgmt^−/−^* NDMA-treated livers compared to WT. Related DAPI quantification provided in Fig. S1E. Scale bar = 20 μm. **d**, Quantification of hepatocyte micronuclei formation by flow cytometry at 2 days post-NDMA exposure. *n* ≥ 10 per group. **e**, Apoptosis quantified by IHC for cleaved caspase-3 (CC3) in *Mgmt*^−/−^ mouse livers. Representative CC3 staining shown in inset. *n* ≥ 2. **f**, Quantification of apoptosis was measured by immunofluorescence of pan-nuclear gH2AX-positive cells for *Mgmt^−/−^* mice. Inset shows representative pan-nuclear gH2AX staining. *n* ≥ 7. **g**, Proliferation measured via flow cytometry for % Ki67-positive cells at 2 days post-exposure. *n* ≥ 10. **h**, PCNA protein quantification by western blot in liver whole-cell lysates at 5 days and 10 weeks post-exposure. Band intensities normalized to TPS and saline WT controls. *n* = 4 per group (2 males and 2 females). Data are presented as mean ± s.e.m. Statistical comparisons performed using one-way ANOVA with Šídák’s multiple comparisons test (**b**,**d**–**h**). Statistical significance: *p < 0.05, **p < 0.01, ***p < 0.001, ****p < 0.0001. NS, not significant. DAPI, 4′,6-diamidino-2-phenylindole.

High levels of DNA damage can cause apoptosis, which prevents propagation of mutant cells. Cleaved caspase-3, a marker for apoptosis, was high in *Mgmt*^−/−^ mice at 1 day and 50 day time points (**Fig. 2e**), but not in WT mice (**Extended Data Fig. 2d**). This programmed cell death was corroborated by prolonged pan-nuclear gH2AX staining across several time points (**Fig. 2f & Extended Data Fig. 2e**). Importantly, apoptosis coincided with regenerative proliferation, as evidenced by an increase in frequency of Ki67-positive cells at 2 days (**Fig. 2g**) and enhanced PCNA expression at 5 days post-exposure (**Fig. 2h**). Proliferation likely raises replication fork encounters with DNA lesions, promoting fork stress and collapse, which may contribute to phenotypic differences in gH2AX persistence.

### Immunoblotting and phosphoproteomics reveal an NDMA-induced DNA damage response

Given the pivotal role of DNA damage responses (DDRs) in determining cell fate, we next examined key DDR proteins and characterized the phosphorylation cascades orchestrating this response^31,32^. Phosphorylation of p53, a canonical ATM substrate, was significantly increased in both WT and *Mgmt*^−/−^ livers at 1 day and 5 days post-exposure and resolved by 10 weeks (**Fig. 3a**). Phospho-p53 induces p21 (CDKN1A) expression, which was markedly stronger in *Mgmt*^−/−^ mice (**Fig. 3b**). Elevated RAD51, indicative of HR activity, was also observed specifically in the *Mgmt*^−/−^ mice (**Fig. 3c**). Induction of p21 was sustained up to 10 weeks in *Mgmt*^−/−^, but not WT mice, demonstrating a persistent DDR that is consistent with sustained gH2AX, described above.

**Fig. 3:**
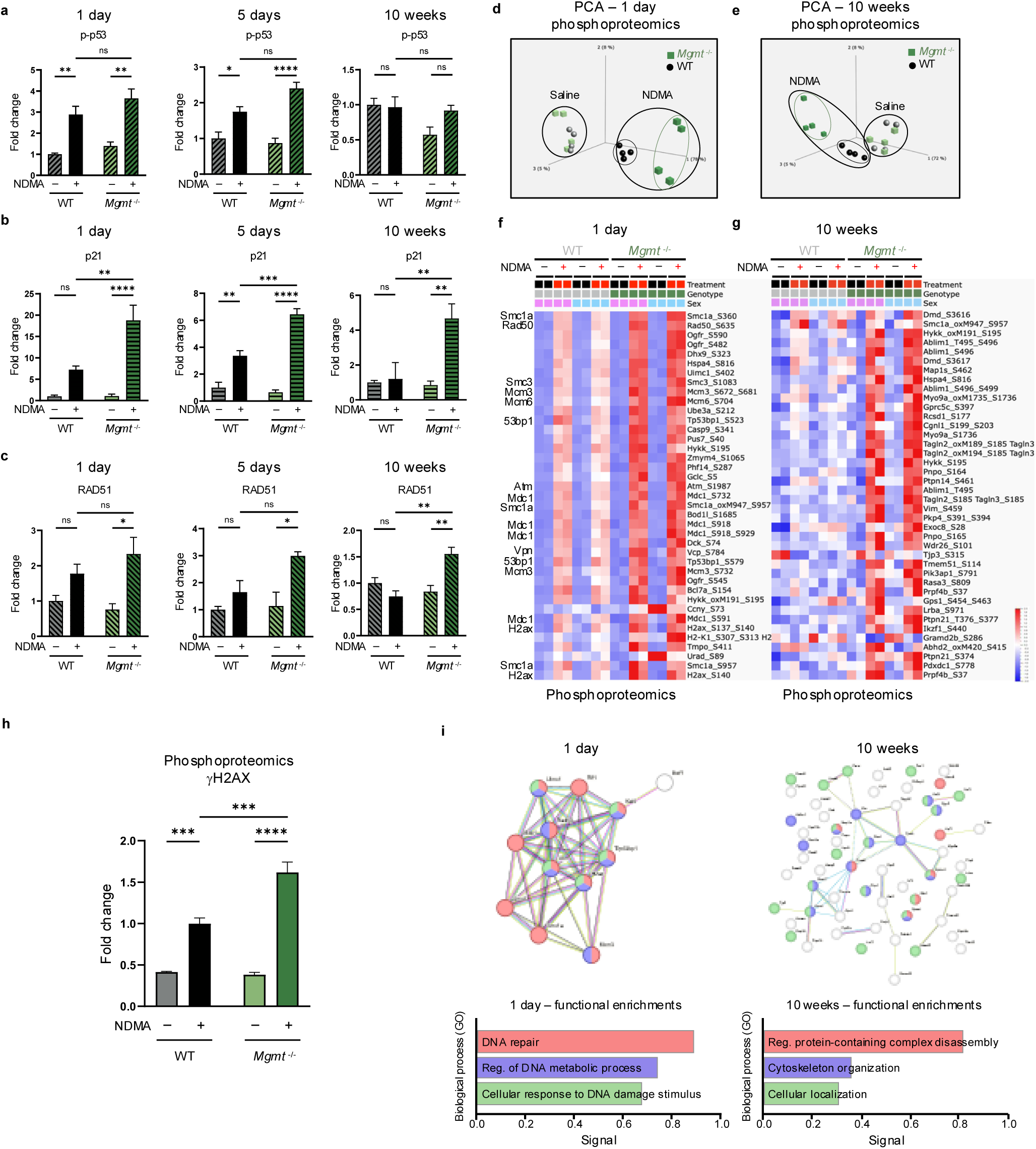
Immunoblotting and phosphoproteomics reveal an NDMA-induced DDR. **a**, Whole-cell liver lysates collected at 1 day, 5 days, and 10 weeks post-exposure were analyzed by western blot for phosho-p53. Band intensities normalized to TPS and saline WT controls. *n* = 4 per group (2 males, 2 females). **b**, Liver lysates from identical timepoints were immunoblotted for p21, normalized as above. *n* = 4 per group (2 males, 2 females). **c**, Liver lysates from identical timepoints were immunoblotted for RAD51, normalized as above. *n* = 4 per group (2 males, 2 females). **d**,**e**, Principal Component Analysis (PCA) of phosphoproteomics at 1 day (**d**) and 10 weeks (**e**) post-exposure displayed distinct sample clustering by treatment group and genotype. *n* = 4 (2 males and 2 females). **f**,**g**, Heatmaps show the most differentially expressed phosphorylated proteins at 1 day (**f**) and 10 weeks (**g**) post-exposure, identified by phosphoproteomics. DNA repair proteins are highlighted in bold (**f**). Protein selection based on multi-group ANOVA by treatment (p < 0.05), with standard deviation < 0.05; 354 total proteins (**f**), 214 total proteins (**g**). **h**, Phosphoproteomic quantification of gH2AX (combined S140 and S137 sites) at 1 day post-exposure. *n* ≥ 2 per group. **i,** STRING database analysis of top-expressed phosphoproteins at 1 day and 10 weeks post-exposure. Functional enrichment at 1 day highlights DNA repair clusters; 10 weeks displays enrichment for non-DNA repair biological processes from gene ontology. Statistical comparisons performed using one-way ANOVA with Šídák’s multiple comparisons test (**a**–**c**,**h**). Data are presented as mean ± s.e.m. Statistical significance: *p < 0.05, **p < 0.01, ***p < 0.001, ****p < 0.0001. NS, not significant.

Phosphoproteomics was performed via mass spectrometry of immunoprecipitated phosphorylated peptides containing the DDR kinase motif (phosphoserine followed by glutamine (pSQ) or phosphothreonine followed by glutamine (pTQ)). Principal component analysis (PCA) demonstrates distinct separation between NDMA-treated and saline control samples at both 1 day and 10 weeks (**Fig. 3d-e**). Only WT and *Mgmt*^−/−^ NDMA-treated groups formed clearly separated clusters, indicating strong shifts in signaling following NDMA exposure. A vast array of proteins were phosphorylated in response to NDMA, almost all of which were more abundant in the *Mgmt*^−/−^ livers (**Fig. 3f-g**). Many of these proteins at 1 day post-exposure, including gH2AX (**Fig. 3h**) and other ATM substrates such as ATM itself (**Fig. 3f & 3i**), are known to be involved in DDR and form clusters in STRING interaction network analysis. Among hundreds of potential ATM targets, there was a strong enrichment for proteins related to HR (including H2AX, Mdc1, Rad50, Smc1a, Uimc1, Smc3, Atm, Vcp, Nbn, and Bod1l) (**Fig. 3f, 3i & Extended Data Fig. 3a**). The fact that *O*^6^MeG adducts trigger activation of machinery relevant to resolution of MMR-induced gaps and broken forks reveals remarkable specificity. These early activation events mediated by phosphorylation may serve as predictive biomarkers for disease or therapeutic responses, as shown previously with the wee1 kinase inhibitor Adavosertib^33^. Although the DDR was robust shortly after exposure, by 10 weeks, this response had shifted to signs of reprogramming of cellular structure, organization, and intracellular communication, with the cytoskeleton and related processes (**Fig. 3g & 3i**).

### NDMA induces persistent transcriptome dysregulation and an IFN response

To advance our understanding of NDMA-induced molecular processes, we studied the transcriptome. PCA of transcriptomic data revealed distinct clustering in NDMA-exposed *Mgmt^−/−^* mice across all early timepoints (**Fig. 4a**). Throughout the first 10 weeks post-exposure, ~1,000 genes remained consistently upregulated in *Mgmt*^−/−^ livers, while WT mice exhibited rapid resolution (**Fig. 4b**). Longitudinal analysis identified 189 shared genes with sustained elevated expression and 55 genes with persistent downregulation in *Mgmt*^−/−^ mice from 1 day all the way to 10 months post-exposure (**Fig. 4c**). In stark contrast, for the WT mice, no genes were identified as being persistently upregulated.

**Fig. 4:**
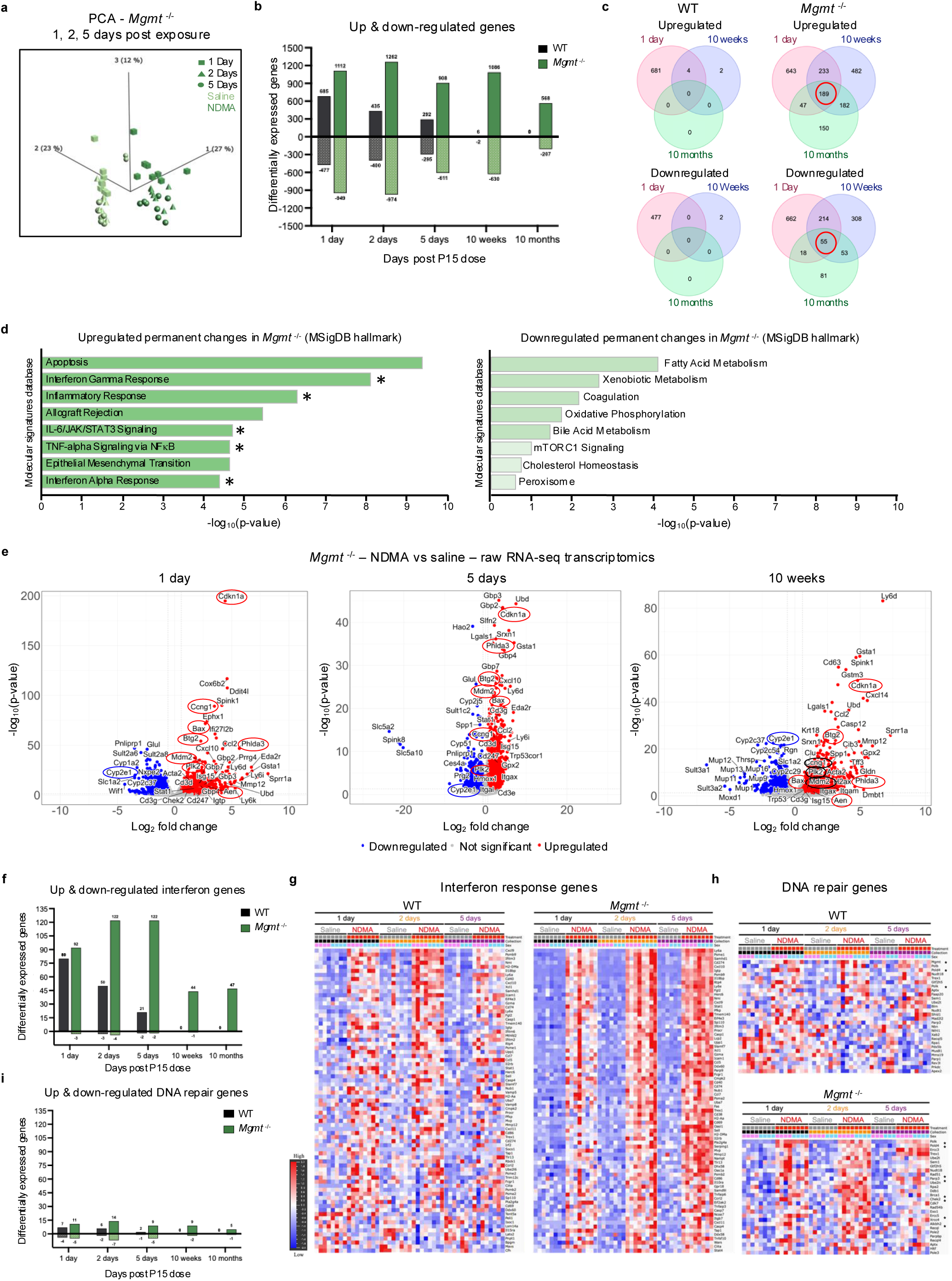
NDMA induces persistent transcriptome dysregulation and an IFN response. **a**, PCA of *Mgmt*^−/−^ liver samples collected at days 1, 2, and 5 post-saline or NDMA exposure shows clear clustering separation by treatment group. RNA was extracted from liver tissue and analyzed by RNA-seq; *n* ≥ 7 per group (males and females combined). **b**, Bar graphs depict the number of differentially expressed RNA transcripts (up- and downregulated) in WT and *Mgmt*^−/−^ liver following NDMA exposure across indicated timepoints. **c**, Venn diagrams compare overlap of significantly up- and downregulated genes across genotypes and timepoints. **d**, Pathway enrichment analysis of persistently up- and downregulated genes (intersected across all timepoints from panel **c**, red circles) in *Mgmt*^−/−^ samples using Enrichr for MSigDB Hallmark gene sets. The top 8 pathways are displayed, ranked by Fisher exact test p-value. Pathways involved in IFN response are marked with an asterisk (*). **e**, Volcano plots display the most differentially expressed genes in *Mgmt*^−/−^ livers comparing NDMA vs saline treatment at 1 day, 5 days, and 10 weeks. Plots show –log_10_ p-value versus log_2_ fold change. Genes from the GSEA Hallmark p53 pathway are circled in red or black; Cyp2e1 downregulation is circled in blue. **f**,**i**, Bar graphs quantify the number of up- and downregulated genes within the pre-curated IFN (**f**) and DNA repair (**i**) gene lists, comparing NDMA effects between WT and *Mgmt*^−/−^ across the time course. **g**,**h**, Heatmaps show expression profiles for curated IFN response genes (**g**) and DNA repair genes (**h**) in response to NDMA treatment in WT and *Mgmt*^−/−^ livers at days 1, 2, and 5 post-exposure. Color scale reflects relative gene expression intensity. Statistical analysis: All comparisons use thresholds of adjusted p-value < 0.05 and absolute log_2_ fold change > 0.58 (**b**– **i**).

To further explore the systems-level responses to NDMA, we performed Enrichr analysis of the Gene Ontology (GO) biological processes and MSigDB pathways. Strikingly, for both WT and *Mgmt*^−/−^ mice, on days 1 and 2 post-exposure, most activated pathways were related to inflammation and the IFN response (**Extended Data Fig. 4a-b**). Furthermore, *Mgmt^−/−^* mice exhibited permanent activation of IFN signaling, IL-6/JAK/STAT3 signaling, TNFα signaling via NF-κB, and apoptosis from 1 day up to 10 months post-exposure (**Fig. 4d**). Concurrently, metabolic pathways were suppressed (**Fig. 4d**), including downregulation of *Cyp2E1* (**Fig. 4e & Extended Data Fig. 4c**), which was further validated at the protein level (**Extended Data Fig. 4d**) and is consistent with chronic liver injury^34–36^. Analysis of the total IFN-related genes (**Fig. 4f & Supplementary Table 1**) and non-hierarchical clustering (**Fig. 4g**) further demonstrate a strong IFN response that never returned to normal in *Mgmt^−/−^* mice (**Extended Data Fig. 4e**). Both non-hierarchical clustering of gene expression data (**Fig. 4h**) and volcano plots (**Fig. 4e & Extended Data Fig. 4c**) reveal a DDR induction, wherein *Mgmt*^−/−^ were more susceptible. Unexpectedly, relatively few genes directly involved in DNA repair were upregulated (**Fig. 4i & Supplementary Table 2**). However, among those induced, many were functionally significant in response to *O*^6^MeG, including HR-related genes (*Brca1*, *Rad51*, & *Ercc5*) and, in addition, *Polk* (**Fig. 4h**), which suppresses *O*^6^MeG-induced toxicity through translesion synthesis^37^. WT mice exhibited similar but attenuated and more transient fluxes in gene expression (**Fig. 4f-i & Extended Data Fig. 4c**). Importantly, both *Gbp3* and *Mgmt* were highly upregulated in WT mice (**Extended Data Fig. 4c, 1 day**). *Gbp3* is stimulated by the IFN response and interacts with STING to enhance expression of *Mgmt* expression^38,39^, likely contributing to the relative resistance of WT mice to disease phenotypes.

NDMA-induced genotoxicity biomarkers were considerably overexpressed, with *Cdkn1a* (*p21*) emerging as one of the most significantly upregulated genes across all timepoints in *Mgmt*^−/−^ mice (**Fig. 4e**). Volcano plots and Gene Set Enrichment Analysis (GSEA) uncovered significant enrichment of the hallmark p53 pathway (**Extended Data Fig. 4f**)^40^. High levels of p21 are thus likely the result of both p53 and IFN pathway activation^41–43^. Taken together, sustained *Cdkn1a* expression and IFN signaling are signatures of the *Mgmt^−/−^* response to NDMA, consistent with a pro-tumorigenic microenvironment observed in this model (**Fig. 1f-g**).

### Upstream regulators of the IFN response are persistently activated in *Mgmt*^−/−^ mice

Analysis using the TRRUST database via Enrichr revealed significant enrichment of specific transcription factors. Notably, IFN-responsive factors showed the strongest signal, with orders-of-magnitude higher enrichment in *Mgmt^−/−^* mice (**Fig. 5a**). QIAGEN Ingenuity Pathway Analysis (IPA) of upstream regulators confirmed a persistent activated IFN response in *Mgmt*^−/−^ mice through 10 months post-exposure (**Fig. 5b**). Specifically, IPA indicated robust induction of both Type I IFN response (IFNα, IFNβ, IFNAR1, IFNAR2, ISGF3, STAT1, STAT2, and IRF9) and Type II IFN response (IFNγ, IFNGR1, JAK1, JAK2, and STAT1 dimer), alongside inhibition of key negative IFN regulators (TREX1, STAT6, IL10RA, USP18, ADAR, SOCS1, and PNPT1) (**Fig. 5b**)^44,45^. In contrast, WT mice exhibited a reduced and more transient IFN response that was entirely absent beyond 5 days post-exposure with no significant genes recorded at 10 weeks or 10 months (**Fig. 5c**). IPA also identified activated DNA damage-related upstream regulators (TP53, ATM, BRCA1, PARP1/2, & CHEK2) in *Mgmt*^−/−^ mice (**Extended Data Fig. 5a**), which diminished over time, unlike the dominant IFN signature.

**Fig. 5:**
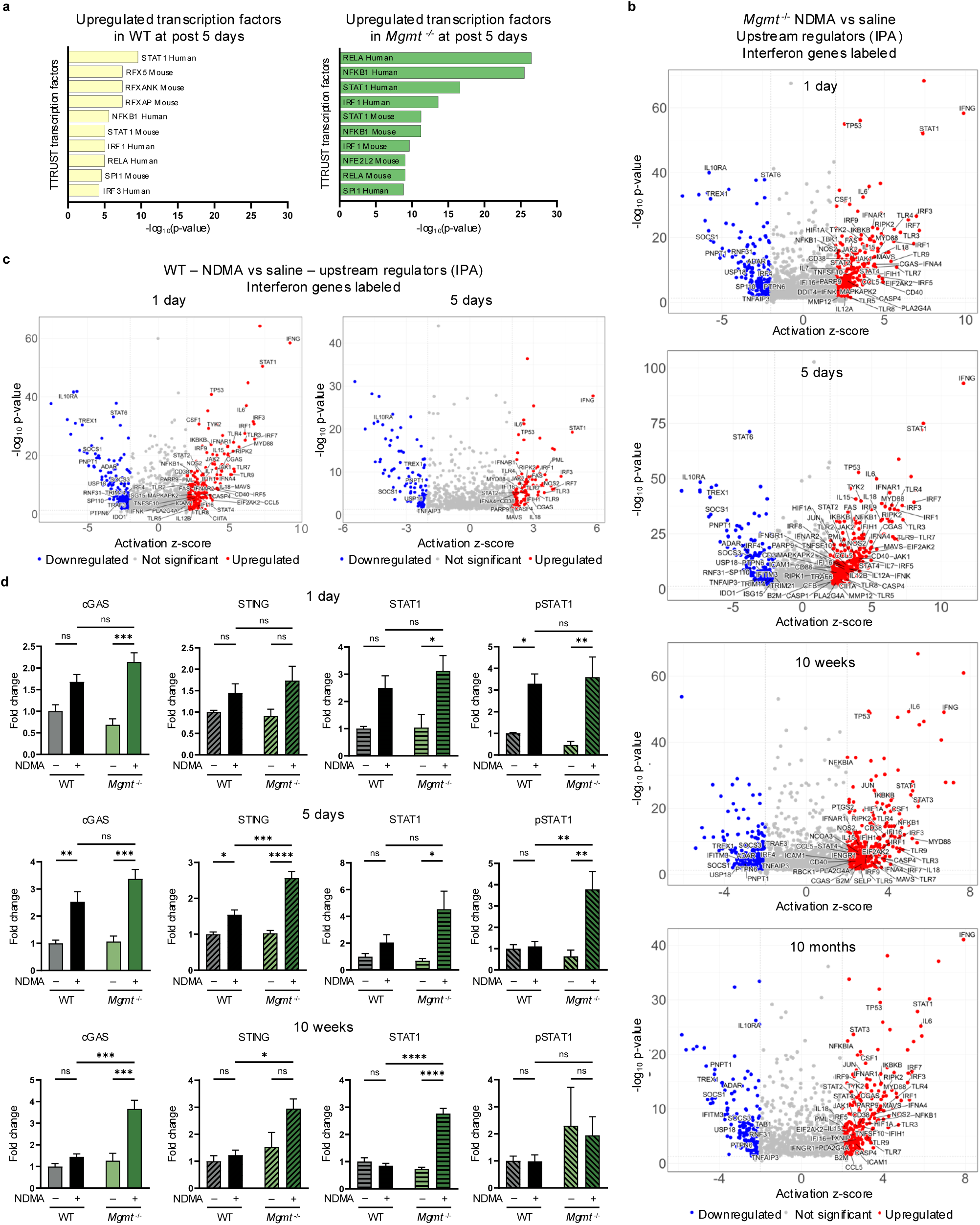
Upstream regulators of the IFN response are persistently activated in *Mgmt*^−/−^ mice. **a**, Upregulated genes from WT and *Mgmt*^−/−^ livers (5 days post-NDMA treatment) were analyzed for transcriptional regulatory network enrichment using the TRRUST (Transcriptional Regulatory Relationships Unravelled by Sentence-based Text-mining) database via Enrichr. Bar graphs display the top 10 enriched transcription factors as ranked by p-value. **b**, Ingenuity Pathway Analysis (IPA) of RNA-sequencing data from *Mgmt*^−/−^ mice identified activated (> 2.0 z-score) and inhibited (< 2.0 z-score) upstream regulators throughout their lifetime with IFN genes labeled in each plot. **c**, IPA of RNA-sequencing data for WT mice identified activated and inhibited upstream regulators only at 1 day and 5 days post-exposure. No genes met cutoff thresholds (same as (B) for 10-week or 10-month data. Genes involved in IFN response are labeled. **d**, IFN pathway protein levels (cGAS, STING, STAT1, pSTAT1) were quantified by western blot from whole-cell liver tissue at 1 day, 5 days, and 10 weeks post-NDMA exposure. Band intensities normalized to TPS and saline WT controls. *n* = 4 per group (2 males, 2 females). Statistical comparisons performed by two-way ANOVA with Šídák’s multiple comparisons test. Data are presented as mean ± s.e.m. Statistical significance: *p < 0.05, **p < 0.01, ***p < 0.001, ****p < 0.0001. NS, not significant. All transcriptomic analyses utilized filtering thresholds of p-adjusted value < 0.05 and absolute log_2_ fold change > 0.58.

IFN regulators, such as STAT1, induce transcription of IFN-stimulated genes, which includes cGAS, STING, as well as STAT1 itself, forming a self-reinforcing feedback loop^46^. Immunoblotting confirmed a persistent upregulation of cGAS, STING, STAT1, and pSTAT1 in *Mgmt*^−/−^ mice with a transient increase in WT mice (**Fig. 5d**), mirroring transcriptional profiles. Phosphorylated IRF7, a master regulator of the Type I IFN response, was also significantly increased exclusively in *Mgmt*^−/−^ mice (**Extended Data Fig. 5b**). These results demonstrate that MGMT deficiency drives a robust and persistent IFN response characterized by concurrent activation of Type I and Type II IFN signaling pathways, a sustained enhancement of effector expression, and key phosphorylation events.

### NDMA induces recombination events indicative of persistent genomic instability

Previous studies have shown that *Mgmt*^−/−^ mice are highly susceptible to point mutations^16,47,48^. Here, we quantified another class of mutations, namely HR-driven sequence-rearrangements. Specifically, we monitored direct repeat recombination at the RaDR locus, which results in EGFP expression that is indicative of large-scale sequence rearrangement mutations (**Fig. 6a-b**)^49^. NDMA exposure strongly induced *de novo* recombination events, peaking 10 days post-exposure (**Fig. 6c & Extended Data Fig. 6a**). While HR frequency was initially comparable between genotypes, *Mgmt*^−/−^ mice exhibited significantly elevated mutations from 30 days onward (**Fig. 6c**). This divergence coincided with reduced liver lobe size in *Mgmt*^−/−^ mice (**Fig. 6d & Extended Data Fig. 6b**), reflecting enhanced pathophysiology (**Fig. 1e & Extended Data Fig. 1c**). We also observed NDMA-induced HR events in the pancreas of *Mgmt*^−/−^ mice (**Extended Data Fig. 6c-e**), suggesting another potentially affected organ and a subject for future exploration. Results thus demonstrate chronic genomic instability and toxicity as central outcomes of NDMA treatment in *Mgmt*^−/−^ mice.

**Fig. 6:**
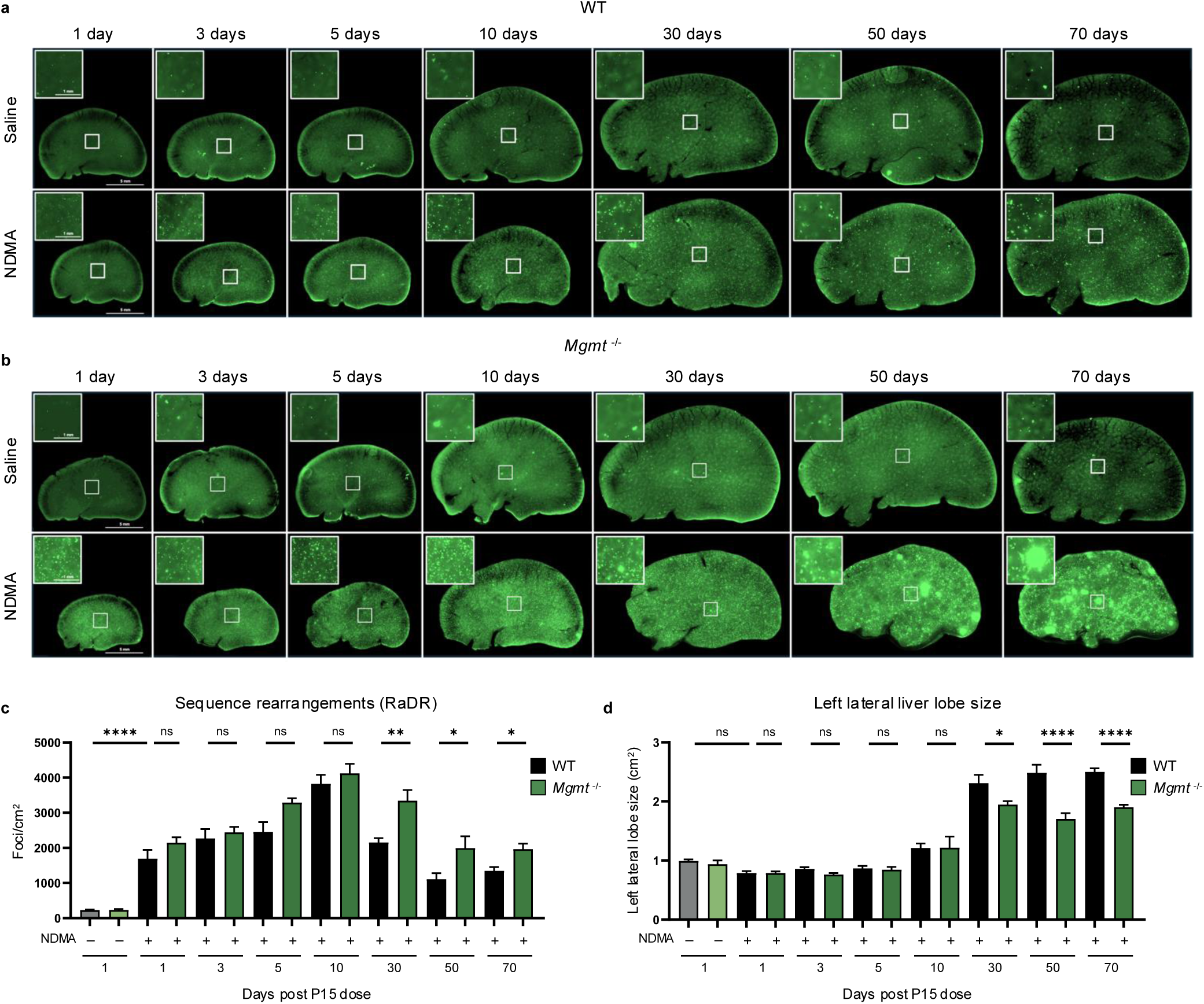
NDMA induces recombination events indicative of persistent genomic instability. **a**, Representative whole-mount fluorescence images of left lateral liver lobes from WT RaDR-GFP mice at multiple timepoints post-NDMA or saline exposure. eGFP-positive foci indicate large-scale sequence rearrangements/recombination events. Insets show higher magnification (scale = 1 mm) of boxed regions (scale = 5 mm at 2x). Images are representative of male mice. **b**, Representative whole-mount fluorescence images of left lateral liver lobes from *Mgmt*^−/−^ RaDR-GFP mice, showing extensive mutation burden and prominent clonal expansion of GFP-positive foci after NDMA treatment. Images are representative of male mice. **c**, Quantitation of sequence rearrangement mutations (eGFP-positive RaDR foci per cm²) in WT (gray/black bars) and *Mgmt*^−/−^ (green bars) livers post-NDMA exposure over time. Foci quantified via machine learning-assisted image analysis. *n* ≥ 10 per group. **d**, Measurements of total left lateral liver lobe area (cm²) in WT (gray/black bars) and *Mgmt*^−/−^ (green bars) mice at various timepoints following NDMA exposure. *n* ≥ 10 per group. Statistical comparisons performed using one-way ANOVA with Šídák’s multiple comparisons test (**c**,**d**). Data are presented as mean ± s.e.m. Statistical significance: *p < 0.05, **p < 0.01, ***p < 0.001, ****p < 0.0001. NS, not significant.

### NDMA induces clonal expansion in *Mgmt^−/−^* livers

An advantage to the RaDR-GFP model is that it is possible to track clonal expansion of recombined EGFP-positive cells. Small clonal outgrowths (>500 pixels) were present in NDMA-exposed *Mgmt^−/−^*mice starting at 30 days post-exposure, while almost none were observed in WT mice (**Fig. 7a**). By 10 weeks, many clones expanded substantially, with a significant proportion exceeding 2000 pixels (**Fig. 6b & 7b**), indicating aggressive clonal expansion for a subpopulation of foci, which may reflect pre-neoplastic changes. These enlarged clones were more frequent in males (**Extended Data Fig. 7a**), which correlates with males’ increased susceptibility to tumor development, raising the possibility that clonal expansion is a driver of sex-specific vulnerability to tumorigenesis (**Extended Data Fig. 1a**).

**Fig. 7:**
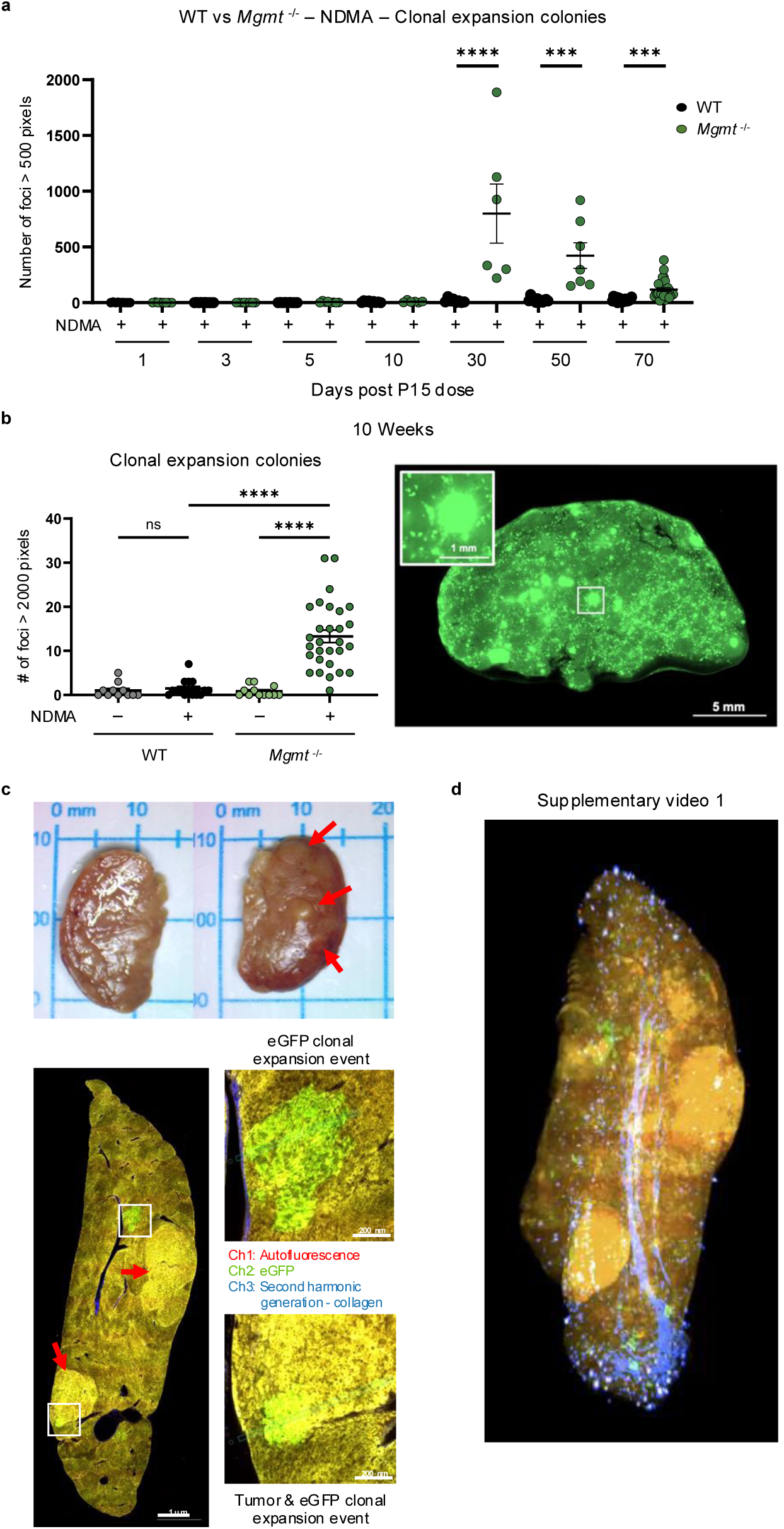
NDMA induces clonal expansion in *Mgmt*^−/−^ livers. **a**, Quantitation of sequence rearrangement mutations that formed clonal expansion colonies in *Mgmt*^−/−^ (green dots) livers starting at 30 days post-NDMA exposure compared to WT (gray/black bars). A single clonal expansion event was defined as RaDR-GFP foci exceeding 500 pixels in area. *n* ≥ 6 per group. **b**, At 10 weeks (70 days) post-NDMA exposure, *Mgmt*^−/−^ livers exhibit significantly larger clonal expansion colonies (RaDR-GFP foci >2000 pixels) compared to WT. A representative eGFP image of a *Mgmt*^−/−^ male mouse liver at 10 weeks post-NDMA exposure highlights clonal expansion colonies. **c**, 2-photon imaging of a ~13-month post NDMA treated *Mgmt*^−/−^ female liver shows both eGFP clonal expansion and tumor expression from autofluorescence (yellow), with corresponding insets displaying high-resolution views. Representative gross liver images highlight visible tumors (red arrows). **d**, 3D reconstruction of sequential 2-photon images from (**c**) visualizes the spatial distribution of eGFP clonal expansion events (green) and tumors (yellow) within a *Mgmt*^−/−^ liver. The vascular network is displayed in blue. See Supplementary Video 1. Statistical comparisons performed using Kruskal-Wallis and Dunn’s test (**a**,**b**). Data are presented as mean ± s.e.m. Statistical significance: *p < 0.05, **p < 0.01, ***p < 0.001, ****p < 0.0001. NS, not significant.

We next performed two-photon microscopy with serial sectioning and deconvolution ^50^ to create 3D images for an entire left liver lobe from a female *Mgmt^−/−^* mouse 1 year post-exposure. This revealed numerous fluorescent RaDR-GFP clonal expansion events (**Fig. 7c-d; Supplementary Video 1 & Extended Data Fig. 7b-c; Supplementary Video 2**). The fact that green fluorescent cells were adjacent to one another verifies that foci are due to clonal expansion, rather than clustering of smaller foci (**Fig. 7d; Supplementary Video 1**). With this method, tumors visible on the surface of the liver were detectable as yellow autofluorescence (**Fig. 7c-d**). An overlap between tumor tissue (yellow) and clonal expansion events (green) indicates clonal expansion within the tumor. Second harmonic generation (blue) visualized vascular networks, revealing spatial relationships among vessels, clones, and tumors. This integrated imaging approach provides unprecedented resolution for mapping clonal dynamics and tumor evolution in intact organs, establishing a transformative platform for studying cancer initiation.

### Spatial transcriptomics of clonal outgrowths and tumors

The RaDR-GFP system provides a unique opportunity to analyze gene expression within clonally expanded fluorescent cell populations. Using 10x Genomics Visium spatial transcriptomics analysis at 10 weeks post-exposure, we found uniform gene expression across liver parenchyma in saline-treated *Mgmt*^−/−^ males (**Fig. 8a**). Examination of EGFP-positive clonally expanded cells in an NDMA-treated liver revealed significant upregulation of *Myc* targets, IFN responses, apoptosis, and p53 pathway genes, indicating activation of multiple stress-response mechanisms suggestive of pre-tumor transformation. Histology uncovered chronic portal hepatitis with biliary hyperplasia, consistent with early pathological changes in this pre-tumor stage (**Fig. 8a, top right**). Additionally, we observed a tumor in a 10-month-old *Mgmt*^−/−^ mouse that exhibited a clearly demarcated boundary when analyzed by H&E staining, spatial transcriptomics, and fluorescence, for which histopathology identified as a hepatocellular adenoma with a few dilated biliary ducts (**Fig. 8b**). This tumor was composed of several enriched pathways that were similar to the 10-week clone (*e.g*., Myc targets and apoptosis). However, there was also upregulation of mTORC1, KRAS, complement, and cholesterol homeostasis signatures, which are known to be crucial for tumor maintenance and progression. Although the RaDR-GFP reporter was not detected within this particular tumor (consistent with a low recombination frequency at the RaDR locus^49^), its stochastic activation nevertheless enabled the identification and molecular analysis of a clonal expansion event, providing a powerful tool for capturing early changes in gene expression that may precede tumorigenesis.

**Figure 8.**
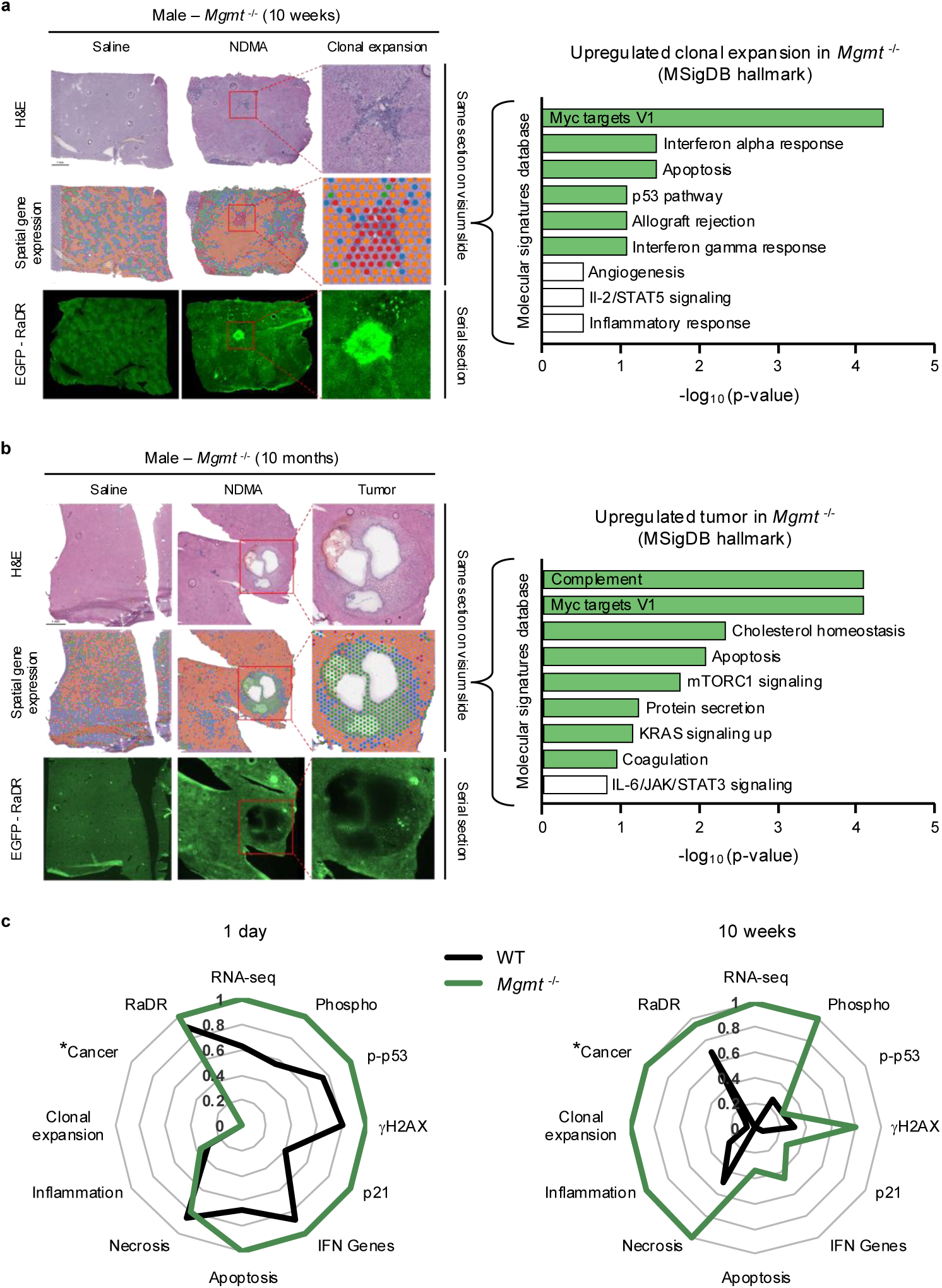
Spatial transcriptomics of clonal outgrowths and tumors. **a**, Spatial transcriptomic profiling was performed on male *Mgmt*^−/−^ liver samples 10 weeks post-treatment using 10x Genomics Visium technology. Representative male *Mgmt*^−/−^ liver samples 10 weeks post-treatment were analyzed. For each sample, the same liver section was used for H&E staining (row 1) and spatial transcriptomics mapping (row 2), while an adjacent serial section was imaged for eGFP fluorescence (row 3). In saline-treated livers, transcriptomic clusters were uniform, but NDMA treatment revealed distinct clustering, accompanied by histological changes including portal hepatitis and biliary hyperplasia, which overlapped with an intensely GFP-positive colony. One featured clonal expansion (~0.5 mm²) was analyzed for differentially expressed RNA transcripts; upregulated genes from this region were enriched for pathways in the MSigDB Hallmark collection, with the top 9 pathways shown. **b**, Spatial transcriptomic profiling of male *Mgmt*^−/−^ livers 10 months post-treatment. Analysis used the same tissue section for H&E staining (row 1) and spatial transcriptomics (row 2); a serial section was imaged for eGFP (row 3). Saline controls continued to show uniform transcriptomic clustering, while NDMA treatment resulted in a hepatocellular adenoma with a few dilated bile ducts. Differentially expressed genes from the tumor region were subjected to MSigDB Hallmark pathway enrichment, with the top 9 pathways shown. **c**, Radar plots summarize key endpoints from this study at 1 day and 10 weeks post-exposure. Medians or means from each data set were normalized to the highest scoring group in that analysis method (e.g., NDMA-treated *Mgmt*^−/−^ mice had the highest tumor burden, so medians of all groups were normalized to the median *Mgmt*^−/−^).

### Data integration across endpoints

To synthesize the main findings, we integrated molecular, cellular, and pathological data from 1 day and 10 weeks following NDMA exposure by generating radar plots (**Fig. 8c**), with each parameter normalized to the highest scoring group for each particular analysis method. Comprehensive analysis clearly shows that, soon after exposure, *Mgmt^−/−^*mice demonstrate heightened DNA damage signaling (phospho-p53, gH2AX), cell cycle arrest (p21), IFN expression, and apoptosis markers. By 10 weeks, these acute molecular responses diminished, while tissue-level pathologies such as necrosis, inflammation, clonal expansion, and early neoplasia (cancer) became more pronounced, reflecting a transition from molecular stress to macroscopic disease. One of the most intriguing findings was the early and persistent responses of transcriptomics, phosphoproteomics and RaDR mutations in *Mgmt^−/−^* mice, suggesting that these measures may serve as predictive biomarkers. This integrated analysis links molecular changes occurring within days of exposure to tissue pathologies observed many weeks later, highlighting their potential to foreshadow subsequent cancer development.

## Discussion

In this study, we investigated how multi-omics responses unfolded following exposure to the carcinogenic compound NDMA early in life, illuminating the relationship between initial DNA damage and subsequent cancer development. By leveraging the *Mgmt*^−/−^ mouse model, we were able to focus specifically on the biological impacts of DNA damage. Within days, the livers of *Mgmt*^−/−^ mice underwent substantial rewiring, with phosphoproteomics showing a robust ATM-driven DDR linked to parallel transcriptional changes. Over two months later, the repercussions of these rapid-onset molecular responses manifested as necrosis, inflammation, and other pathologies. Pathway analysis revealed several upregulated processes, wherein the IFN response was the most significant and enduring. To our knowledge, this study is among the first to demonstrate a direct link between a methylating agent and the induction of a robust IFN response. The observation that DNA damage elicited a sustained IFN response, from 1 day to 10 months after exposure, highlights the potential for a burst of DNA damage to initiate a persistent immune response that promotes liver damage. Thus, acute exposure to an alkylating agent at a young age can lead to long-term tissue pathology and disease.

Comprehensive pathway analyses highlighted both Type I and Type II IFN responses as potential drivers of hepatocellular microenvironment reprogramming in response to NDMA. These responses were likely driven by cGAS-STING or other cytosolic DNA sensors^51–53^. The observation that over a hundred genes related to the IFN response were upregulated along with phosphorylation of IRF7 (a key mediator of the IFN response) demonstrated a mechanistic link between DNA methylation damage and innate immune activation^54^. IFN responses have been previously implicated in DNA damage responses^24,55–58^, and our study demonstrated their robust and sustained activation *in vivo* in mice exposed transiently to DNA damage.

However, the IFN response to NDMA exposure can have opposing effects. Early on, there were DNA-protective mechanisms, such as induction of *p21* (which mediates cell cycle arrest), *Isg15* (which enhances p53 activity^59^), and in WT Mice, *Gbp3* (which promotes *Mgmt* expression^38,39^). While these responses are beneficial in the short term, a persistent IFN response could promote chronic pro-inflammatory responses, for example via activation of NF-kB^60^, which was observed through IPA at 10 months post-exposure in *Mgmt*^−/−^ mice. Furthermore, IPA also identified several Toll-like receptors (TLRs) and their adaptor protein MYD88, which are upstream regulators of the IFN response^61^. Importantly, TLR4-KO mice become resistant to inflammation and hepatic fibrosis following exposure to the ethylating agent diethylnitrosamine^25^. It remains to be established whether diminished TLR activity offers similar protection against methylation-induced IFN activation and ensuing liver toxicity.

MGMT silencing is common in tumors that are sensitive to temozolomide (TMZ), whereas MGMT-proficient tumors tend to be resistant^62–64^. Clinical trials have explored the use of IFN agonists to promote immunogenic tumor cell death, in some cases in combination with traditional chemotherapies^65,66^. However, for methylating agents like TMZ, IFN agonists may have limited efficacy in treating MGMT-silenced tumors, as we demonstrated that methylation damage alone induced an IFN response in *Mgmt*^−/−^ mice. In contrast, IFN agonists could be beneficial in MGMT-proficient tumors, since MGMT expression could suppress TMZ-induced IFN responses. Thus, patient stratification by a tumor’s MGMT status may be critical when combining methylating agents with IFN agonists, which carry their own risks.

In our model, both WT and *Mgmt*^−/−^ mice initially exhibited similar intensity across numerous phenotypes, including levels of gH2AX, recombination frequency, IFN activation, and markers of pathophysiology, although phosphoproteomic and transcriptomic responses were strikingly different between WT and *Mgmt*^−/−^. This immediate hyperactivation of signal transduction and gene expression, together with persistent IFN signaling, may underlie the predictive power for methylation-driven carcinogenesis. Analysis of temporal dynamics revealed a marked divergence by 10 weeks, with all phenotypes significantly stronger in *Mgmt*^−/−^ mice. Thus, disease prediction based on these phenotypes relied primarily on delayed rather than immediate responses.

Our multi-omics approach enabled an integrated look at homologous recombination (HR). Phosphoproteomics showed that some of the most frequently phosphorylated proteins are directly involved in HR. Notably, HR-related genes were also upregulated at the transcriptional level, and IPA further identified activated genes in this pathway through its integration of extensive upstream regulator networks. Consistent with these observations, RaDR-GFP quantification revealed significant induction of HR-mediated sequence rearrangement mutations. Taken together, this work demonstrated that *O*^6^MeG is recombinogenic *in vivo*, which is consistent with *in vitro* studies showing HR to be a critical defense against *O*^6^MeG-induced DSBs^67,68^.

We validated that the RaDR-GFP model detects clonal expansion by demonstrating shared spatially constrained gene expression patterns, as well as adjacent fluorescent cells within clonal outgrowths when visualized in 3D. As such, RaDR-GFP offers a visual model for detecting this early stage of tumorigenesis. By peering into clonal expansion events via spatial transcriptomics, we revealed localized IFN activation and concurrent immune modulation. Oncogenic pathways were also upregulated within clonal events, including *Myc*, *p53*, *mTORC1*, and *Kras*. These may be driven by *O*^6^MeG-induced mutations, which have been observed in MGMT-silenced malignancies such as hepatobiliary, lung, and gastric cancers^62,63^. Further, longitudinal analysis identified conserved transcriptomic dysregulation in proliferative signaling, connecting early clonal expansions to late tumorigenesis.

For these studies, we treated mice with two doses of NDMA as this regimen had been shown previously to induce tumors^12,27^. However, a weakness of this approach is that responses to the first dose overlap with responses to the second dose. These conditions likely obscured detailed response kinetics, such as RaDR-GFP recombination events. Studies of a single dose are ongoing and will refine our understanding of the precise timing of responses.

Collectively, our findings demonstrated that an early and transient alkylation insult in an MGMT-deficient context elicited immediate activation of DDRs as well as unrelenting IFN signaling. Such persistent IFN activation may have driven toxicity that enabled clonal expansion and microenvironment remodeling to promote tumor development. These results implicate rewiring of phosphorylation events, disruption of transcription, and sustained IFN-driven processes as critical events early in carcinogenesis and highlight their potential utility as predictive biomarkers and targets for disease mitigation.

## Resource availability

### Lead contact

Further information and requests for resources and reagents should be directed to and will be fulfilled by the Lead Contact, Bevin Engelward.

### Materials availability

This study did not generate new unique reagents.

### Data and code availability

The accession number for the mass spectrometry proteomics data reported in this paper is ProteomeXchange Consortium via the PRIDE (Perez-Riverol et al., 2019) partner repository: PXD021142. The RaDR foci counting algorithm is available on Github at https://github.com/dushanw/RPN_RCN_roiExtractor or at figshare at https://figshare.com/articles/software/RPN_RCN_roiExtractor/29581370?file=56318189

All raw data files can found at the MIT Superfund Research Program Fairdom hub at https://fairdomhub.org/projects/221

## Methods

### Animals

All experimental mice were on a C57BL/6 genetic background. *RaDR*^R/R^;*gpt*^g/g^ (WT) mice were created by crossing RaDR-GFP mice (B6.129S4(Cg)-*Gt(ROSA)26Sor^tm1(CAG-EGFP*)Bpeng)^*/J)^49^ with *gpt* delta mice^69^. *RaDR*^R/R^;*gpt*^g/g^;*Mgmt*^−/−^ mice were generated by crossing *RaDR*^R/R^;*gpt*^g/g^ and *Mgmt*^−/−^ mice (described previously^48^). The mice were maintained in AAALAC-certified animal care facilities and provided standard food and water *ad libitum*. Mice were housed in static microisolator cages with autoclaved hardwood chip bedding and a nestlet. Macroenvironmental conditions included temperature maintenance at 70 ± 2°F, 30–70% humidity, and a 12:12 h light:dark cycle. Mice were free of the following murine pathogens: ecto- and endoparasites, mouse parvovirus, mouse hepatitis virus, mouse rotavirus, minute virus of mice, ectromelia, sendai virus, pneumonia virus of mice, reovirus, theilovirus, lymphocytic choriomeningitis virus, *mycoplasma pulmonis*, *filobacterium rodentium*, polyoma virus and mouse adenovirus. Pathogen status was verified by full serology testing using sentinel mice every six months, and with an abbreviated serology panel every two months. All animal procedures were performed according to the NIH Guide for the Care and Use of Laboratory Animals and protocols approved by the Massachusetts Institute of Technology Committee on Animal Care.

### NDMA treatment

NDMA was produced in-house, verified through NMR and mass spectrometry (MS) and stored at −20°C as previously described^12^. Litters of mice were designated for saline or NDMA treatment at birth. A total dose of 10.5 mg/kg NDMA diluted in saline was administered intraperitoneally over two separate injections, according to Dass et al^27^. First, 3.5 mg/kg NDMA in saline (10 μL volume) was given at 8 days of age (P8), and the remaining 7 mg/kg (20 μL volume) was given at 15 days of age (P15) (**Fig. 1a**). Control mice were sham treated at the same time points with equivalent volumes of saline.

Mice were euthanized by asphyxiation with carbon dioxide according to AVMA guidelines and necropsied. Whenever possible, equal numbers of males and females of each genotype and treatment group were analyzed for each endpoint. When data were collected from an excess number of males or females for an endpoint, individual data points were selected by random number generator for exclusion from analysis.

### Animal analysis

Most analyses combined data from male and female mice unless specifically mentioned in the figure legends. Analyses included mice from separate litters whenever possible to reduce litter-based differences. Each mouse represented an experimental unit (*n =* 1). Numbers of mice and treatment timepoints included are described in the figure legends. An equal number of male and female mice from each cohort were used when possible. For some endpoints with excess mice of one sex, individuals were randomly excluded from analysis to balance group sizes.

### DNA adduct analysis

#### Isolation of DNA from frozen liver tissue, DNA hydrolysis and solid-phase extraction

Adduct levels were quantified for the livers of mice treated with NDMA at days P8 and P15, whereas saline-treated mice received treatment only at P15. DNA was isolated from 100 mg of frozen liver tissue using the Qiagen 100/G genomic tip (Qiagen) following the procedure recommended for genomic DNA preparation. Briefly, liver tissue was homogenized in buffer QBT in the presence of RNase and proteinase K and incubated overnight at room temperature. The solution was loaded onto a Qiagen 100/G tip, washed with buffer QC, and retained DNA was eluted with buffer QF. DNA was precipitated with isopropanol and recovered by spooling onto a glass rod, which was washed with 70% ethanol, briefly dried and then dissolved in 500 μL of purified water. The amount of recovered DNA was determined via UV absorption using a NanoDrop One C spectrometer (ThermoScientific, Waltham, MA, USA).

DNA (100-125 μg) containing 10 μg of methylated ^15^N DNA, which served as an internal standard, was hydrolyzed by addition of 50 μL formic acid and heating at 95°C for 1 h. Hydrolyzed DNA samples containing isotopically labeled standards were loaded onto Strata X SC cartridges (100 mg; Phenomenex, Torrance, CA), which had been prewashed with acetonitrile followed by water. The cartridges were then washed with 15 mL of 0.1 N hydrochloric acid. The retained compounds were eluted with 1.5 mL of a solution containing 5% ammonium hydroxide, 10% ethanol, and water, and vacuum centrifuged to dryness at room temperature. Samples were reconstituted in 100 μL of 0.1 N HCl and transferred to glass vials for LC/QQQ MS analysis.

#### LC/QQQ MS analysis of DNA adducts

LC/ESI-MS/MS analysis was performed on an Agilent 6410 Triple Quadrupole mass spectrometer interfaced with an Agilent 110 series HPLC (Agilent Technologies, Palo Alto, CA). Samples (25 μL injected) were resolved on a 4.6 x 150 mm, 3.5 mm Eclipse Plus C18 column (Agilent Technologies, Palo Alto, CA) eluted at a flow rate of 0.5 mL/min. The column was eluted with a step gradient of 4 mM ammonium acetate containing 2% acetonitrile (solvent A) to 100% acetonitrile/0.0001% acetic acid (solvent B) over a period of 30 min followed by column re-equilibration. QQQ MS analysis was performed in the positive ion mode using N_2_ as the nebulizing/drying gas. Capillary voltage was 4.0 kV and temperature was set at 350°C. Selective reaction monitoring (SRM) was used for quantitative analysis of methylated purines with the collision energy set at 25 for methylated guanines. Nitrogen was used as the collision gas. The following transitions were monitored: *m/z* 166 [M+H^+^] → m/z 149 for m6G; *m/z* 171 [M+H^+^] → *m/z* 153 for *O*^6^-methyl-^15^N-guanine.

### Phosphoproteomics

Methods used were consistent with prior work^12^. Briefly, 1 day and 10 weeks after the second injection, liver samples were flash frozen in liquid nitrogen, and stored at −80°C. 2 males and 2 females for each genotype and treatment were analyzed. Liver tissues were homogenized in ice-cold 8 M urea (Sigma) with three 10 second pulses. Proteins were processed, digested and desalted as described previously^70^. Lyophilized peptide aliquots of 400 mg (of starting protein) were labeled with TMT10-plex labeling kits (Thermo Fisher). Phosphopeptides were enriched by immunoprecipitation (IP) followed by Fe-NTA-based immobilized metal affinity chromatography (IMAC). TMT-labeled samples were resuspended in IP buffer (100 mM Tris-HCl, 1% Nonidet P-40, pH 7.4) and incubated overnight with PTMScan Phospho-ATM/ATR Substrate Motif kit (Cell Signaling Technology). Peptides were eluted twice, each with 25 mL of 0.2% trifluoroacetic acid (TFA) for 10 minutes at room temperature followed by a secondary Fe-NTA-based IMAC to remove non-specifically retained non-phosphopeptides.

High-Select Fe-NTA enrichment kit (Pierce) was used according to manufacturer’s protocol with following modifications. After washing the Fe-NTA spin columns, beads were resuspended in 25 mL of binding washing buffer. Eluates from IP were incubated with Fe-NTA beads for 30 minutes. Peptides were eluted twice with 20 mL of elution buffer into a 1.7-mL microcentrifuge tube. Eluates were dried in SpeedVac until 1–5 mL of sample remained. Samples were resuspended in 10 mL of 5% acetonitrile/0.1% formic acid and loaded directly onto an in-house packed analytical capillary column (50 mm ID x 10 cm) packed with 5-mm C18 beads (YMC gel, ODS-AQ, AQ12S05).

Liquid chromatography tandem mass spectrometry (LC-MS/MS) of phosphopeptides and crude lysate analysis was carried out on an Agilent 1260 LC coupled to a Q Exactive HF-X mass spectrometer (Thermo Fisher) as described previously^70^. Raw mass spectra data files were processed with Proteome Discoverer version 2.2 (Thermo Fisher) and searched against the mouse SwissProt database using Mascot version 2.4 (Matrix Science). TMT reporter quantification was extracted using Proteome Discoverer. MS/MS spectra were searched with the following settings: a) mass tolerance of 10 ppm for precursor ions; b) 20 mmu for fragmentations; c) fixed modification for cysteine carbamidomethylation; d) TMT-labeled lysine; e) TMT-labeled peptide N-termini; and f) dynamic modifications for methionine oxidation and phosphorylation of serine, threonine and tyrosine. Peptide spectrum matches (PSMs) were filtered according to following parameters: rank = 1, search engine rank = 1, mascot ion score > 20, isolation interference < 30%, average TMT signal > 1000. Peptides with missing values across any channel were filtered out. Phosphorylation sites were localized using ptmRS module^71^ on Proteome Discoverer. PSMs with > 95% localization probability were included for further analysis. Only peptides containing ‘SQ’ or ‘TQ’ sequence motif were included for final analysis. Peptide quantification was normalized with relative median values obtained from crude peptide analysis. Further data analysis was performed in Python (version 3.6) and MATLAB (R2016a).

#### STRING analysis

Protein selection based on multi-group ANOVA by treatment (p < 0.05), with standard deviation < 0.05 were used to select 354 total proteins for 1 day analysis and 214 total proteins for 10 weeks analysis. These proteins were subjected to STRING database analysis to identify predicted protein-protein interactions. Gene ontology biological processes were selected for cluster identification and comparisons.

### RaDR analysis

The methods used were consistent with prior work^12^. Briefly, livers were collected from mice across the time course starting at 1 day after the second dose injection to 10 weeks post-exposure. Freshly excised livers were held on ice in 0.01% trypsin inhibitor (Boston BioProducts) in PBS prior to imaging (**Extended Data Fig. 8a**). The entire left lobe of the liver was secured between a glass slide and a coverslip. The dorsal surface of each liver was then imaged with a Nikon Eclipse Ti2 scanning microscope on the 2x objective in the FITC: EGFP channel using an Andor Zyla 4.2 camera and NIS Elements software (v5.11.02). Emission wavelength was 535 nm and all images were captured under identical exposure settings across samples.

A user-trained two-stage machine learning algorithm was used to identify and enumerate fluorescent foci within intact tissue (available on Github at https://github.com/dushanw/RPN_RCN_roiExtractor) and (available on figshare at https://figshare.com/articles/software/RPN_RCN_roiExtractor/29581370?file=56318189). Similar to our prior studies^72^, we first identified potential regions (termed Region Proposals) that might contain foci and then classified them into True Foci or False Regions. Two deep convolutional neural networks (DCNNs) were trained using 10 manually annotated images for both tasks. The first Region Proposal Network segmented the foci-like regions, which were used to extract the corresponding image patches (Region Proposals). These Region Proposals were then fed to a second DCNN (Proposal Classifier Network) that classified them into either True Foci or False Regions. True Foci locations were then listed and counted to get the final foci count. The 10 manually annotated training images comprised samples from each genotype and treatment group. All liver images were analyzed by the machine learning program based on parameters developed from training data. The number of fluorescent foci was normalized to the area of the liver in the image. The left lateral liver lobe size was extracted from this output. For clonal expansion analysis we based the average size of a single fluorescent focus at 10 pixels using ImageJ (Fiji) (ImageJ v2.9.0). Fluorescent clusters were then measured by size and frequency for foci >500 pixels and >2000 pixels, which were considered clonal expansion events.

Recombination events can happen anytime in the course of liver development. The vast majority of events happen late in development. Livers having undergone recombination early in development are identifiable by EGFP morphology (**Extended Data Fig. 8b**). Livers harboring these early events (7/327) were not used for further analysis.

#### RaDR clonal expansion

For clonal expansion analysis, a custom ImageJ (v1.54p) macro was used to quantify the size of fluorescent foci. Processed images underwent Gaussian blur, thresholding, and watershed segmentation to define regions of interest. Foci greater than 500 pixels (0.00525 mm²) or 2000 pixels (0.021 mm²) were counted as a clonal expansion event based on visual confirmation. The number of clonal expansion events per left liver lobe were quantified.

### Histological analysis

The methods used were consistent with prior work^12^. Briefly, sections of liver were fixed in 10% buffered formalin, embedded in paraffin, and sectioned at 4 µm thickness using a microtome, followed by hematoxylin and eosin (H&E) staining. Liver sections from 4 males and 4 females of each group (1 day, 10 weeks and 10 months post-treatment) were scored by a board-certified veterinary pathologist blinded to sample identity. Specific lesions were graded with a numerical score from 0 to 4; where 0 = normal, 1 = minimal, 2 = mild, 3 = moderate, and 4 = severe. The following hepatic lesions were graded: inflammation, hepatocellular degeneration, hepatocellular necrosis, nuclear enlargement (karyomegaly), hepatic lipidosis, extramedullary hematopoiesis, kupffer cell hyperplasia, Ito cell hyperplasia, bile duct hyperplasia/dysplasia, and fibrosis^73^. Lesions of the liver scored as present (1) or absent (0) included hemorrhage and neoplasia. A total inflammation score for each liver was generated by combining individual scores of portal, midzonal, and centrilobular inflammation from each section^74^. Foci of altered hepatocytes were classified based on morphologic criteria reported by Thoolen and colleagues^73^. Sections were examined using an Olympus BX41 microscope attached with an iKona digital camera and photographed.

### Immunofluorescence

The methods used were consistent with prior work^12^. Briefly, formalin-fixed tissue sections (4 µm) (*n =* 4 males and 4 females from each treatment group and time point) were deparaffinized in xylenes, rehydrated, and subjected to heat-induced antigen retrieval with Dako Target Retrieval Solution (Agilent, #S1699). Sections were blocked with 5% bovine serum albumin, 0.4% Triton X-100 and 5% goat serum for 1 hour at room temperature, then incubated with an antibody for γH2AX (1:200; Cell Signaling Technologies (S139 – 20E3 Rabbit mAb #9718) in 1% BSA and 0.5% Tween 20 in PBS overnight at 4°C. Sections were then washed and incubated with a secondary antibody conjugated to an AlexaFluor 488 probe (1:400; Invitrogen goat anti-rabbit IgG #A1108 (Lot#2382186)) for 1 hour at room temperature. Nuclei were counterstained with ProLong^TM^ Gold AntiFade with DNA Stain DAPI (Invitrogen, #P36931). Stained tissue sections were imaged at 20x under DAPI and FITC filters using a Nikon Eclipse Ti2 microscope and an Andor Zyla 4.2 camera using NIS-Elements AR software (v5.11.02), with two independent regions randomly selected from each slide.

A custom ImageJ (Fiji, v1.54p) macro performed background subtraction, channel-specific thresholding (gH2AX), and nuclear region-of-interest (ROI) generation via DAPI segmentation. Mean fluorescence intensity for gH2AX was quantified within individual nuclei and averaged across two images per experimental replicate. Cells containing a mean pixel intensity greater than 1000 background subtraction were counted as γH2AX foci-positive. Cells with greater than 50% gH2AX staining were considered pan-nuclear and apoptotic. Segmented particles were counted in ImageJ for an approximate number of nuclei in the image. Data were collected for at least 1000 nuclei from each image. Average nucleus size was calculated by pixels using ImageJ.

### Caspase IHC

The methods used were consistent with prior work^12^. Formalin-fixed tissue sections (4 µm) from 4 males and 4 females from each treatment group and time point (same livers as those analyzed for γH2AX and histopathology) were deparaffinized in xylenes, rehydrated, and subjected to HIER in Buffer H, pH 8.8 (Thermo Scientific). After cooling, endogenous peroxidase was blocked with Peroxidazed 1 (Biocare Medical) for 5 minutes and slides were blocked in Background Sniper (Biocare Medical) for 15 minutes. Slides were then incubated with cleaved caspase-3 primary antibody (Cell Signaling Technology - Asp175 (D3E9) Rabbit mAb #9579) for 60 minutes at room temperature, washed, and incubated with Rabbit-on-Rodent HRP Polymer (Biocare Medical) for 30 minutes at room temperature. Slides were washed and incubated with DAB (Betazoid DAB Chromagen kit, Biocare Medical), washed, and counterstained with hematoxylin. Stained slides were scanned with a Leica Aperio AT2 slide scanning microscope and analyzed with QuPath software^75^. After blinding filenames, 4 equal-sized, randomly selected, representative, independent regions of tissue were annotated for number of nuclei and number of apoptotic events. The percentage of apoptotic events for each animal was calculated from the total number of apoptotic hepatocytes and total number of nuclei from four representative 400x fields.

### Micronucleus assay

The methods used were consistent with prior work^12^. Briefly, mice (*n =* 4 males, 4 females of each group) were euthanized 48 hours after the second NDMA injection and livers were collected into 1 mL of ice-cold Liver Preservation Buffer, packed on ice, and shipped overnight to Litron Laboratories. Upon receipt at Litron, each liver was removed from the Liver Preservation Buffer, patted dry, placed into a separate flask containing 10 mL of Liver Rinse Solution, and processed as described previously^76^ using Prototype In Vivo MicroFlow® PLUS ML Kits. Samples were analyzed with a FACSCanto II flow cytometer equipped with 488 and 633 nM excitation (BD Biosciences, San Jose, CA). Instrumentation settings and data acquisition/analysis were controlled with FACSDivaTM software v6.1.3 (BD Biosciences). SYTOX Green-associated fluorescence emissions were collected in the FITC channel (530/30 band-pass filter), and anti-Ki67-eFluoR 660-associated fluorescence emissions were collected in the APC channel (660/20 band-pass filter). The flow cytometry gating strategy for hepatocyte micronucleus (MNHEP) scoring required events to fall within each of three regions and one histogram marker before they were scored as nuclei or micronuclei. The incidence of flow cytometry-scored MNHEP is expressed as frequency percent (no. micronuclei/no. nuclei x 100) of 20,000 SYTOX Green positive nuclei per specimen. Simultaneous with micronucleus assessments, an experimental index of hepatocyte proliferation was collected based on gating for Ki67-positive nuclei.

MNHEP microscopy was performed by adding SYTOX® Green-stained cells to acridine orange-coated slides, then imaged on an Olympus BH-2 microscope with a 40x objective as previously described^76^.

### Tumor analysis

Gross surface lesions on the entire liver were recorded at necropsy. Distinct macroscopic tumors were enumerated; multifocal to coalescing tumorous regions were counted as one.

### RNA sequencing

#### RNA extraction

Total RNA was extracted from ≤30 mg mouse liver tissue stored in RNAlater at −20°C using the RNeasy Plus Mini Kit (Qiagen, cat. no. 74134). Tissue was homogenized in Buffer RLT Plus with β-mercaptoethanol (10 μL/ml) using a BeadBug bead-milling homogenizer at 400 rpm for two 30-second intervals. Lysates were centrifuged, and supernatant was processed through gDNA Eliminator and RNeasy spin columns according to the manufacturer’s protocol. RNA was eluted in 30–50 μL RNase-free water and quantified by NanoDrop spectrophotometry (Thermo Scientific).

#### Library preparation and sequencing

3’ Digital gene expression libraries were created as previously described^77,78^. RNA samples were cleaned using SPRI beads (1.8x) and quantified and quality assessed using an Advanced Analytical Fragment Analyzer. 10 ng of total RNA was used for library preparation. 3’DGE-custom primers 3V6NEXT-bmc#1-24 were added to a final concentration of 1 μM. (5’-/5Biosg/ACACTCTTTCCCTACACGACGCTCTTCCGATCT[BC6]N10T30VN-3’ where 5Biosg = 5’ biotin, [BC6] = 6 bp barcode specific to each sample/well, N10 = Unique Molecular Identifiers, Integrated DNA technologies), to generate two subpools of 24 samples each. After addition of the oligonucleotides, Maxima H Minus RT was added per manufacturer’s recommendations with Template-Switching oligo 5V6NEXT (10uM, [5V6NEXT : 5’-iCiGiCACACTCTTTCCCTACACGACGCrGrGrG-3’ where iC: iso-dC, iG: iso-dG, rG: RNA G]) followed by incubation at 42°C for 90 minutes and inactivation at 80°C for 10 minutes. Following the template switching reaction, cDNA from 24 wells containing unique well identifiers were pooled together and cleaned using RNA Ampure beads at 1.0x. cDNA was eluted with 17 μL of water followed by digestion with Exonuclease I at 37°C for 30 minutes and inactivated at 8 °C for 20 minutes.

Second strand synthesis and PCR amplification was done by adding the Advantage 2 Polymerase Mix (Clontech) and the SINGV6 primer (10 pmol, Integrated DNA Technologies 5’-/5Biosg/ACACTCTTTCCCTACACGACGC-3’) directly to the exonuclease reaction. 8 cycles of PCR were performed followed by clean up using regular SPRI beads at 0.6x and eluted with 20 μL of EB. Successful amplification of cDNA was confirmed using the Fragment Analyzer. Illumina libraries were then produced using standard Nextera tagmentation substituting P5NEXTPT5-bmc primer (25 μM, Integrated DNA Technologies, (5’-AATGATACGGCGACCACCGAGATCTACACTCTTTCCCTACACGACGCTCTTCCG*A*T* C*T*-3’ where * = phosphorothioate bonds) in place of the normal N500 primer. Final libraries were cleaned using SPRI beads at 0.7x and quantified using the Fragment Analyzer and qPCR before being loaded for paired-end sequencing using the Illumina NextSeq500 in paired-end mode (28F,8I,50R nt reads).

#### Data processing and analysis

Sequencing data were aligned to the mouse genome (GRCm38) using STAR aligner, and gene expression was quantified using the ESAT pipeline. Differential expression was performed using DESeq2(median-of-ratios normalization)^79^. Log₂ fold changes were shrunk using lfcShrink to shrink dispersion^80^. Metadata including sample sex, treatment, genotype, and timepoint were incorporated using custom R scripts. Volcano plots were generated from DESeq2 results (log₂ fold changes and p-values). QIAGEN Ingenuity Pathway Analysis (IPA) was used to identify upstream regulators from the differentially expressed genes according to the manufacturer’s instructions using the built-in algorithms.

#### Pathway enrichment

Genes with adjusted p-value < 0.05 and log_2_ fold change > 0.58 were analyzed for pathway enrichment using Enrichr and the Molecular Signature Data Base (MSigDB) Hallmark 2020 mouse gene set or Gene Ontology Biological Process or Transcriptional Regulatory Relationships Unravelled by Sentence-based Text-mining (TRRUST) or Gene Set Enrichment Analysis (GSEA). Pathways with p-value < 0.05 were included for visualization.

After RNA-seq was completed Spliced Transcripts Alignment to a Reference (STAR) was performed to mouse genome reference mm10.

Expression analysis of the digital expression (DGE) libraries targeting transcript ends was performed using the End Sequencing Analysis Toolkit (ESAT). ESAT takes a set of alignment files (SAM or BAM) with genome alignment coordinates, a file containing transcript coordinates (BED or text file) and outputs read counts for each transcript provided. ESAT can be found here on GitHub: (src/java/umms/esatJar/esat.latest.jar).

The package DESeq2 was used for differential expression analysis and can be found here: (http://www.bioconductor.org/help/workflows/rnaseqGene/). DESeq2 uses negative binomial generalized linear models which uses estimates of dispersion and logarithmic fold changes to incorporate data-driven prior distributions.

Lastly, normalization was performed.

### Cluster identification and annotation

Principal component analysis (PCA) was performed on the scaled data to reduce the dimensions, with number of components chosen based on a cumulative proportion (accumulated amount of explained variance) of 95.

A comprehensive analysis of literature yielded a curated list of 337 genes related to the IFN response (Supplementary Table 1). The DNA repair gene list was published by Wood et al.^81–84^ (Supplementary Table 2). Qlucore software was used to generate heatmaps and PCA plots.

### Spatial transcriptomics

Freshly frozen samples were processed following 10x Genomics (Visium) protocols. In brief, fresh frozen tissues embedded in OCT compound (Tissue-Tek) were sectioned at a thickness of 10 µm, the tissue was placed in a Visium glass slide capture frame of 6.5 mm^2^ regions, incubated with methanol at −20°C and then washed with isopropanol. H&E staining was applied, followed by imaging on an Aperio AT2 slide scanner. RNA release was achieved by permeabilization, spatial barcodes were added by reverse transcription and DNA was synthesized and amplified for library preparation. Each capture region has an approximate 5000 spot array, each with a diameter of 50 µm. Sequencing was performed on a NextSeq500 with targeted read depths of at least 100 million reads per sample. A serial section was imaged for RaDR foci and clonal expansion for EGFP to overlay with spatial transcriptomics and H&E. H&E was analyzed and scored by a board-certified veterinary pathologist blinded to sample identity.

### Western blot

Antibodies used for western blots are listed in Supplementary Table 3. Flash-frozen liver samples were homogenized in ice-cold lysis buffer (20 mM Tris pH 7.5, 150 mM NaCl, 1% Triton X-100) supplemented with protease/phosphatase inhibitors. Lysates were sonicated, centrifuged (14,000 × g, 15 min, 4°C), and protein concentrations determined using a BCA assay. Equal protein amounts (40–50 µg) were resolved on 12% SDS-PAGE gels and transferred to nitrocellulose membranes (LI-COR Odyssey) overnight at 30 V (4°C). Membranes were stained with LI-COR Revert 700 Total Protein Stain for normalization and imaged (LI-COR CLx).

For target detection, membranes were blocked in LI-COR Intercept Blocking Buffer (TBS) and incubated overnight at 4°C with primary antibodies diluted in blocking buffer containing 0.2% Tween-20. After washing with TBST, membranes were incubated with LI-COR IR Dye secondary antibodies for 1 hour at room temperature. Signals were visualized on a LI-COR CLx Imager and quantified using Image Studio Lite (v5.2), normalized to total protein. Stripping and reprobing were performed as needed using LI-COR NewBlot buffers. Representative blot images are provided in Extended Data Figures 9 and 10.

### Serial two-photon tomography

Left lateral liver lobes were harvested, drop-fixed in 4% paraformaldehyde in PBS for 24 hrs, then transferred to 4°C 0.1% Sodium Azide in PBS. The livers were then embedded in agarose and prepared for imaging by TissueVision on the TissueCyte 1000 system (TissueVision Inc, Newton, MA; www.tissuevision.com) equipped with a Nikon 16x water immersion objective and a Ti:Sapph laser (MaiTai BB 990G, Spectra Physics). This system uses Serial Two-Photon Plus (STP^2^) Tomography of consecutive sections at 50 μm apart^50^. High-resolution data were collected to construct second harmonic generation (SHG) images arising from endogenous contrast of collagen fibers (920 nm excitation, 20x 1.0 NA objective, emission filter bandpass 442−478 nm). To image deep within the sample, the microscope is equipped with a microtome on the stage so that the sample can be sectioned following collection of each 3D image stack. The imaging resolution and sectioning parameters were as follows: imaging volume = 1.4 μm/pixel *x*,*y*; 2 μm *z*; 25 optical sections for total imaging depth of 50 μm and physical sections = 100 μm thick; 25 sections for a total tissue depth of 2.5 mm. Fluorescence emission was divided into three channels for collection: Channel 1: > 560 nm (Red), Channel 2: 500–560 nm (Green - eGFP), and Channel 3: 442–478 nm (Blue - SHG).

### Quantification and statistical analyses

Statistical analyses were performed using the GraphPad Prism software with the following test: tumor multiplicity (Kruskal-Wallis test with Dunn’s multiple comparisons test for sex-pooled data and ordinary one-way ANOVA with Šídák’s multiple comparisons test for sex-seperated data), RaDR foci (ordinary one-way ANOVA with Šídák’s multiple comparisons test), RaDR clonal expansion (Kruskal-Wallis test with Dunn’s multiple comparisons test or Mann-Whitney test), caspase staining (ordinary one-way ANOVA with Šídák’s multiple comparisons test), histopathology scores (ordinary one-way ANOVA with Tukey’s multiple comparisons test), body weight (ordinary one-way ANOVA with Šídák’s multiple comparisons test), phosphoproteome (two-tailed Student’s t test corrected for multiple hypothesis testing based on Benjamini-Hochberg false discovery rate (FDR) correction), flow cytometry (micronuclei and Ki67) (ordinary one-way ANOVA with Šídák’s multiple comparisons test), γH2AX (ordinary one-way ANOVA with Šídák’s multiple comparisons test), and western blot (two-way ANOVA with Šídák’s multiple comparisons test). A p-value was considered significant if less than 0.05.

## Acknowledgements

This research was supported by the NIEHS Superfund Research Program Grant (P42 ES027707), NIEHS Core Center Grant (P30-ES002109), NIH Training Grant (T32-ES007020), and the Anonymous Fund for Climate Action. The MIT Division of Comparative Medicine assisted in creating animal strains and providing histological services. We thank Caroline Atkinson and Ed Clark of the MIT Division of Comparative Medicine for their contributions to histological processing. We also thank Kathleen Cormier of the MIT Koch Institute Histology Facility for her histology services with spatial transcriptomic processing. We thank the MIT Biomicro Center for providing RNA sequencing services and the MIT Bioinformatics and computing Facility for assisting with RNA sequencing analysis. We gratefully acknowledge Dushan Wadduwage for providing the machine learning code for RaDR foci analysis, as described in Kay et al.^12^. Schematics were created with BioRender.com.

## Contributions

L.J.P. and B.P.E. conceived the ideas, with support from J.E.K., J.J.C., and L.D.S. L.J.P designed the study, wrote the manuscript, and prepared the figures with substantial support from B.P.E. and additional assistance from J.J.C., L.B.V., and M.N. L.J.P. interpreted the data with major contributions from B.P.E. Mouse planning, treatment, collections, and RaDR acquisition were primarily performed by L.J.P., J.E.K., and J.J.C. J.J.C. performed RaDR foci analysis with support from L.B.V., as well as RaDR clonal expansion analysis with custom ImageJ code created by L.B.V. S.E.C. contributed to histopathology analysis and interpretation. Immunofluorescence staining and imaging was performed by L.J.P., M.R.S., and I.S.N. Immunofluorescence analysis with custom ImageJ code was created and performed by L.B.V., with major contributions from E.A.K. Caspase-3 analysis was performed by E.A.K. using QuPath. M.N. performed western blot experiments with significant support from L.B.V. Western blots were analyzed by L.B.V. and L.J.P. with support from M.N. R.G.C. and E.M. performed DNA adduct analysis with support from J.M.E. and L.J.P. N.A.O. performed RNA extractions. S.S.L. performed RNA sequencing. A.C.M. wrote batch code for RNA sequencing differential expression comparisons with support from D.M. N.A.O. and L.J.P. analyzed RNA sequencing data with significant support from B.P.E. and D.M. M.N. produced volcano plots using custom R scripts, with support from D.M. E.A.K. produced heatmaps using Qlucore and curated interferon gene list with support of L.J.P. IPA was performed by L.J.P. Phophoproteomics was performed and analyzed by I.N.K. with support from F.M.W and L.J.P. Micronucleas assay and analysis was performed by D.K.T., S.L.A., and S.D.D. at Litron Labs. 2-photon microscopy was performed by T.R. at TissueVision. NDMA was provided by R.G.C. and J.M.E. All authors reviewed and approved the final manuscript.

## Ethics declarations

The authors declare no competing interests.

## Extended Data Figures

**Extended Data Fig. 1:**
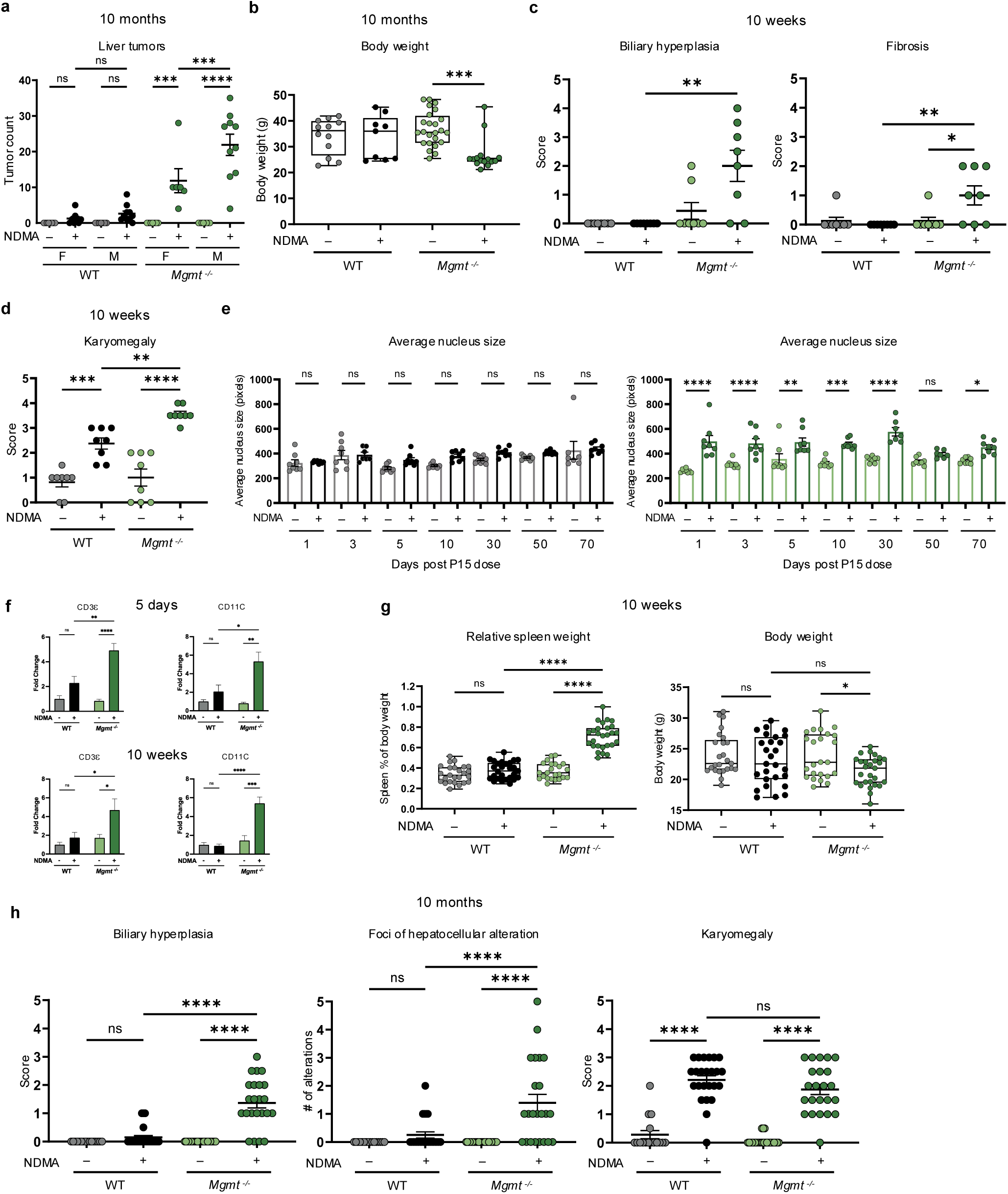
MGMT deficiency exacerbates NDMA-induced liver injury and tumorigenesis. **a**, Liver tumor counts at 10 months post-treatment in WT and *Mgmt*^−/−^ mice, grouped by sex. *n* ≥ 6 per group. **b**, Mouse body weight at 10 months post-treatment (males and females combined). *n* ≥ 9. Box plots indicate median, upper and lower quartiles, and whiskers showing maximum/minimum values. **c**, Histopathological scores for liver morphologies, biliary hyperplasia and fibrosis, in WT and *Mgmt*^−/−^ mice at 10 weeks post-treatment, assessed by H&E staining (sexes combined). *n* ≥ 8. **d**, Histopathological scores of karyomegaly in WT and *Mgmt*^−/−^ mice at 10 weeks post-treatment, assessed by H&E staining (sexes combined). *n* ≥ 8. **e**, Quantification of mean nuclear size in liver cells by DAPI fluorescence. WT saline-(gray) and NDMA-treated (black). *Mgmt*^−/−^ saline-(light green) and NDMA-treated (dark green). See Fig. 2C for images. *n* ≥ 7. **f**, Protein levels were measured via western blot to support T-cell activation (CD3ε & CD11C) in *Mgmt*^−/−^ and WT livers at 5 days and 10 weeks post-exposure. *n =* 4 (2 males, 2 females). **g**, Relative spleen weight (to body weight) and absolute body weight at 10 weeks post-treatment. *n* ≥ 23. Box plots as described in (B). **h**, Histopathological scores for biliary hyperplasia and karyomegaly and number of foci hepatocellular alterations in WT and *Mgmt*^−/−^ mice at 10 months post-treatment, assessed by H&E staining (sexes combined). *n* ≥ 16. Data are presented as mean ± s.e.m. Statistical comparisons performed using one-way ANOVA with Šídák’s multiple comparisons test (**a**,**b**,**e**–**g**) and Kruskal-Wallis and Dunn’s test (**c**,**d**,**h**). Statistical significance: *p < 0.05, **p < 0.01, ***p < 0.001, ****p < 0.0001. NS, not significant.

**Extended Data Fig. 2:**
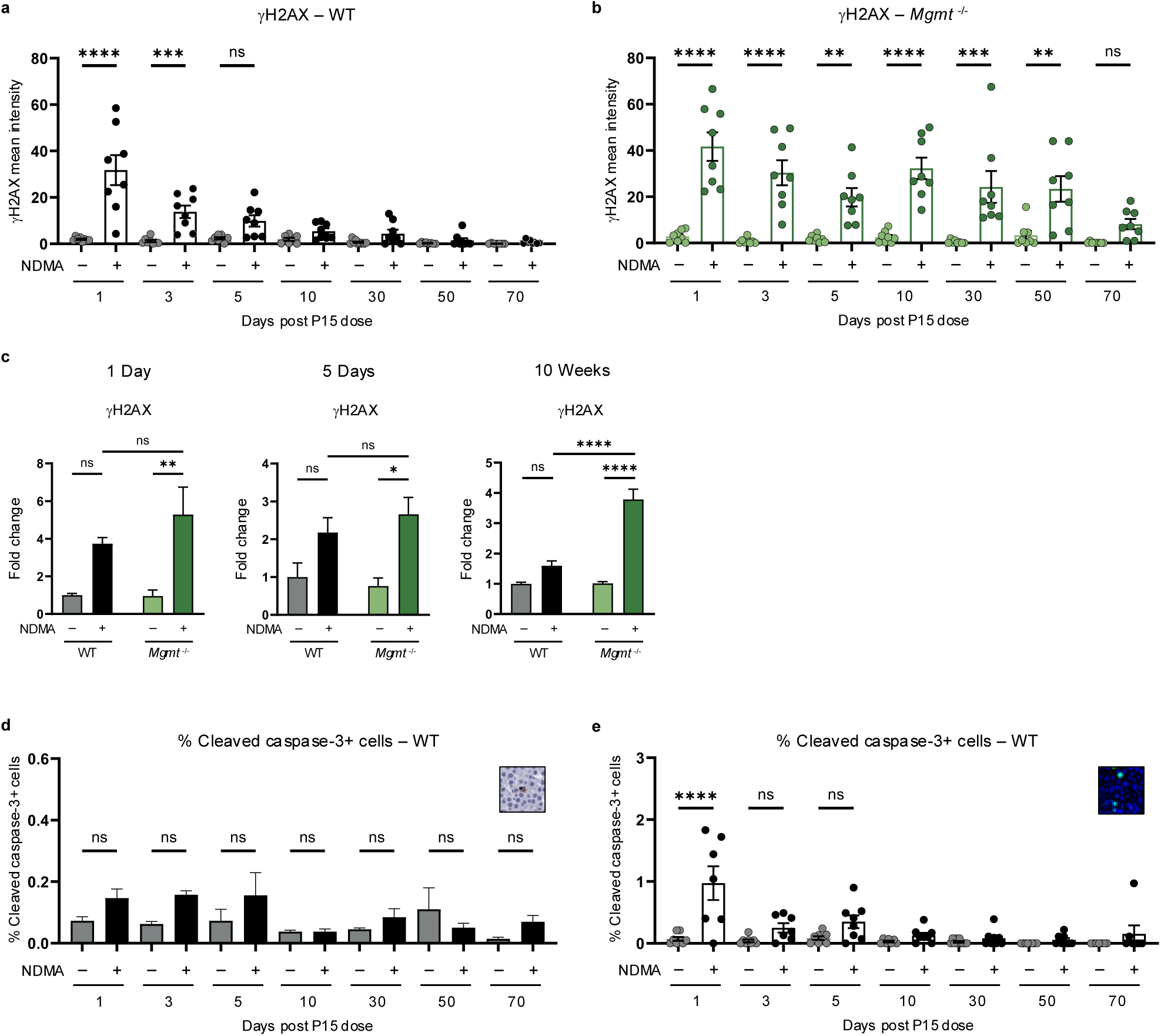
MGMT suppresses DNA damage retention, toxicity, and compensatory proliferation. **a**, Quantification of mean gH2AX fluorescence intensity per nucleus in WT livers, comparing saline-treated (gray) and NDMA-treated (black) groups. *n* ≥ 7 per group. **b**, Quantification of mean gH2AX fluorescence intensity per nucleus in *Mgmt*^−/−^ livers, comparing saline-treated (light green) and NDMA-treated (dark green) groups. *n* ≥ 7 per group. **c**, Western blot analysis of gH2AX in whole-cell liver lysates at 1 day, 5 days, and 10 weeks post-exposure. Band intensities normalized to TPS and saline WT controls. *n* = 4 per group (2 males, 2 females). **d**, Quantification of apoptosis via cleaved caspase-3 positive cells as a percentage of total cells in WT livers at indicated timepoints, measured by IHC. Inset shows representative cleaved caspase-3 staining. *n* ≥ 7. **e**, Quantification of apoptosis via immunofluorescence of pan-nuclear gH2AX-positive cells as a percentage of total cells in WT livers. Inset provides representative pan-nuclear gH2AX staining. *n* ≥ 7. Data are presented as mean ± s.e.m. Statistical comparisons performed using one-way ANOVA with Šídák’s multiple comparisons test (**a**,**b**,**d**,**e**,**f**) and two-way ANOVA (**c**). Statistical significance: *p < 0.05, **p < 0.01, ***p < 0.001, ****p < 0.0001. NS, not significant.

**Extended Data Fig. 3:**
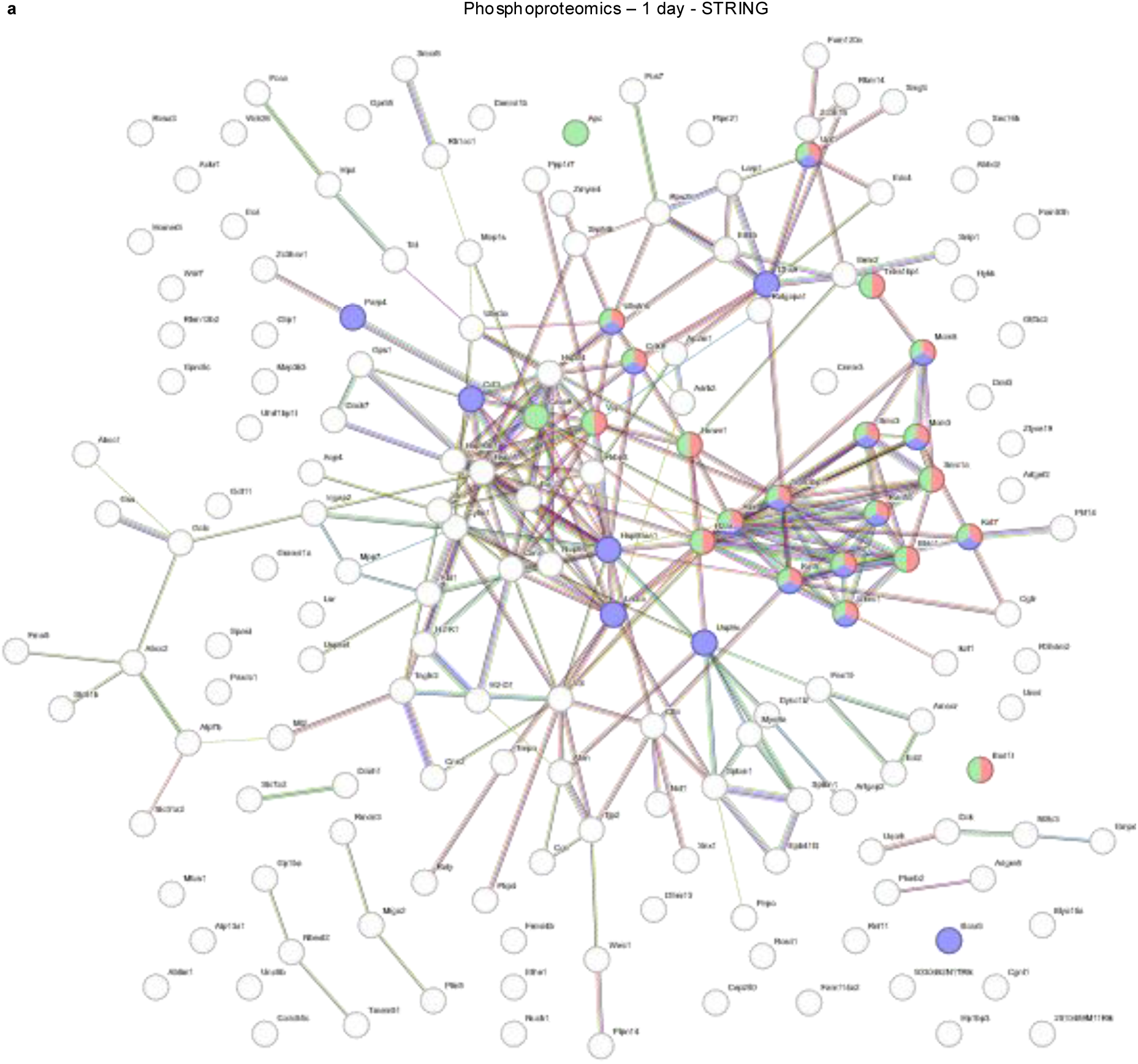
Immunoblotting and phosphoproteomics reveal an NDMA-induced DDR. **a**, STRING database analysis of top-expressed phosphoproteins at 1 day post-exposure. DNA repair clusters as shown in Fig. 3i from biological processes in gene ontology are shown here.

**Extended Data Fig. 4:**
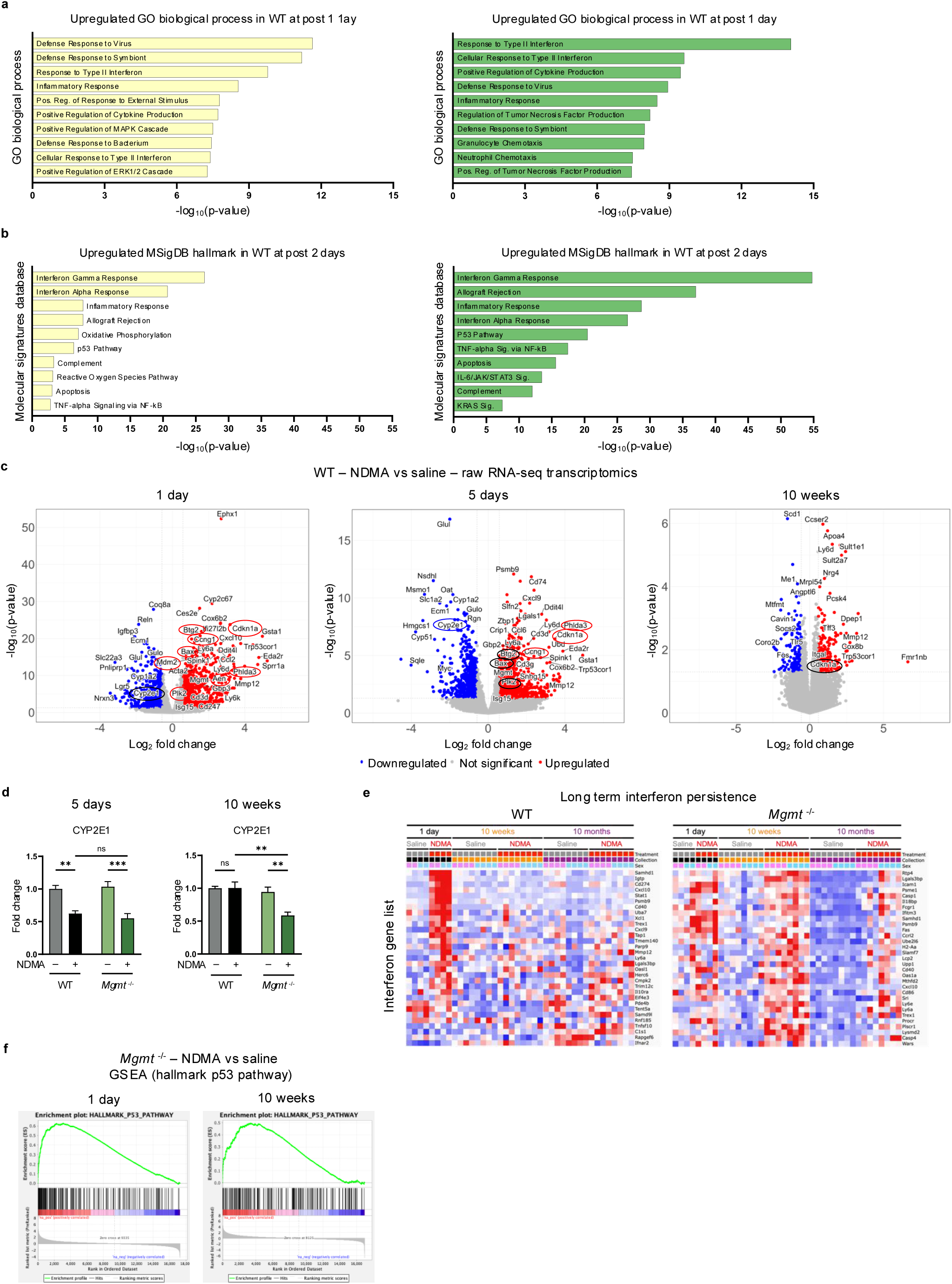
NDMA induces persistent transcriptome dysregulation and an IFN response. **a**, Upregulated genes from NDMA-treated WT and *Mgmt*^−/−^ liver samples (1 day post-exposure) were analyzed using Enrichr for Gene Ontology Biological Process enrichment. The top 10 enriched pathways are shown, ranked by Fisher exact test p-values. **b**, Upregulated genes from NDMA-treated WT and *Mgmt*^−/−^ liver samples (2 days post-exposure) were analyzed using Enrichr for MSigDB Hallmark gene set enrichment. The top 10 pathways are displayed, ranked by Fisher exact test p-values. **c**, Volcano plots present differentially expressed genes in WT liver (NDMA vs saline) across selected timepoints (1 day, 5 days, and 10 weeks), with −log10 p-value plotted against log_2_ fold change. Genes circled in red belong to the GSEA Hallmark p53 pathway data set; Cyp2e1 is circled in blue. Differential expression was determined using an adjusted p-value cutoff of < 0.05 and absolute log_2_ fold change cutoff of > 0.58. **d**, CYP2E1 protein abundance was assessed by western blot in liver samples at 5 days and 10 weeks post-exposure. Band intensities normalized to TPS and saline WT controls. *n* = 4 per group (2 males, 2 females). Statistical comparisons (two-way ANOVA with Šídák’s multiple comparisons. Data are presented as mean ± s.e.m. Statistical significance: *p < 0.05, **p < 0.01, ***p < 0.001, ****p < 0.0001. NS, not significant. **e**, Heatmaps display expression of a curated interferon response gene list in WT and *Mgmt*^−/−^ livers at 1 day, 10 weeks, and 10 months post-exposure, demonstrating persistence of IFN signaling in *Mgmt*^−/−^ samples. **f**, Gene Set Enrichment Analysis (GSEA) on *Mgmt*^−/−^ liver at 1 day and 10 weeks post-NDMA exposure highlights enrichment of the Hallmark p53 pathway compared to saline controls.

**Extended Data Fig. 5:**
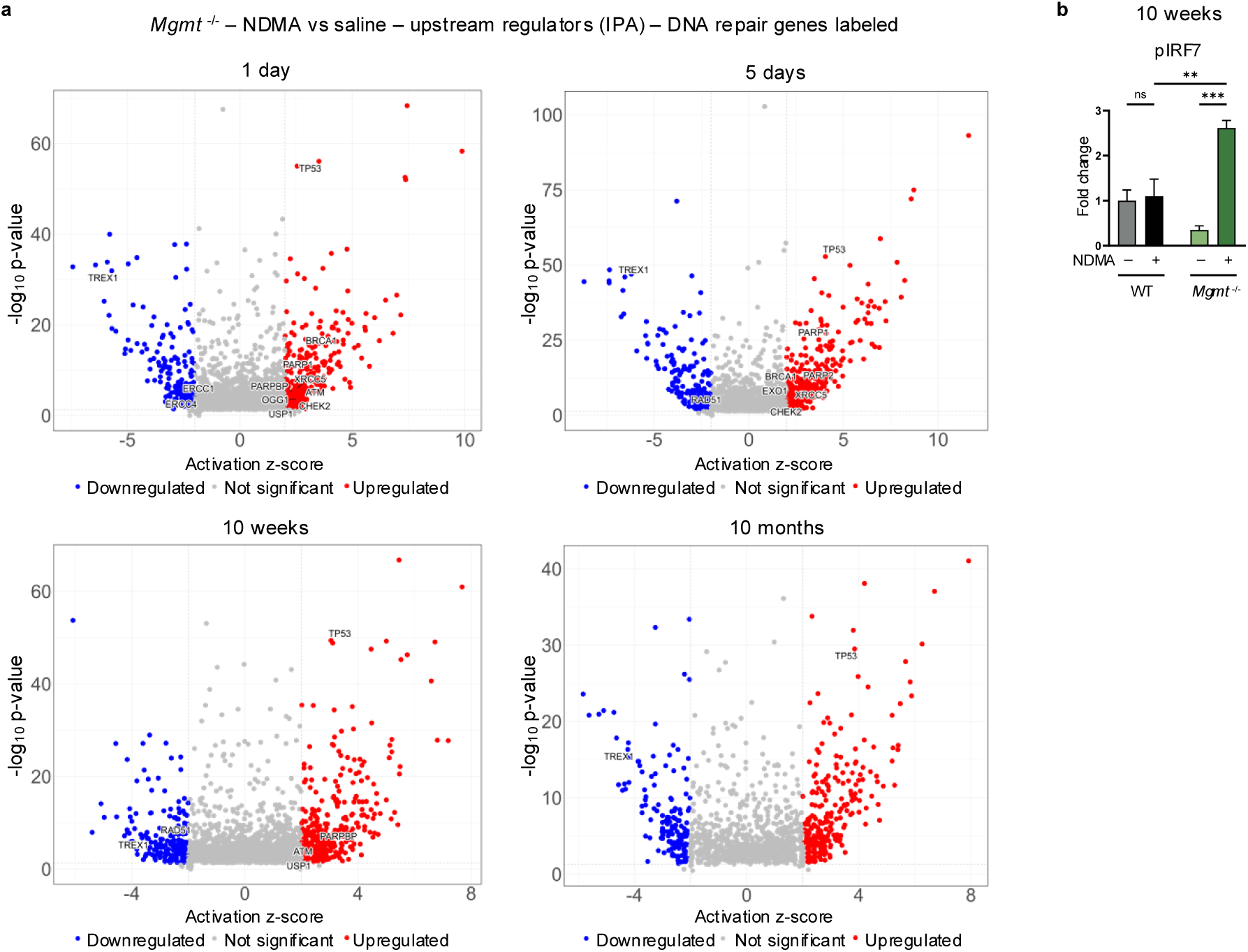
Upstream regulators of the IFN response are persistently activated in *Mgmt*^−/−^ mice. **a**, Ingenuity Pathway Analysis (IPA) of RNA-sequencing data from *Mgmt*^−/−^ mice identified a few activated (> 2.0 z-score) and inhibited (< 2.0 z-score) upstream regulators throughout their lifetime with pertaining to DNA repair genes (labeled in each plot). Analysis included transcripts with p-adjusted values < 0.05 and log_2_ fold change > 0.58. **b**, Phosphorylated IRF7 (pIRF7) protein levels were measured via western blot at 10 weeks post-exposure. Band intensities normalized to TPS and saline WT controls. *n* = 4 per group (2 males, 2 females). Statistical comparisons performed by two-way ANOVA with Šídák’s multiple comparisons test. Data are presented as mean ± s.e.m. Statistical significance: *p < 0.05, **p < 0.01, ***p < 0.001, ****p < 0.0001. NS, not significant.

**Extended Data Fig. 6:**
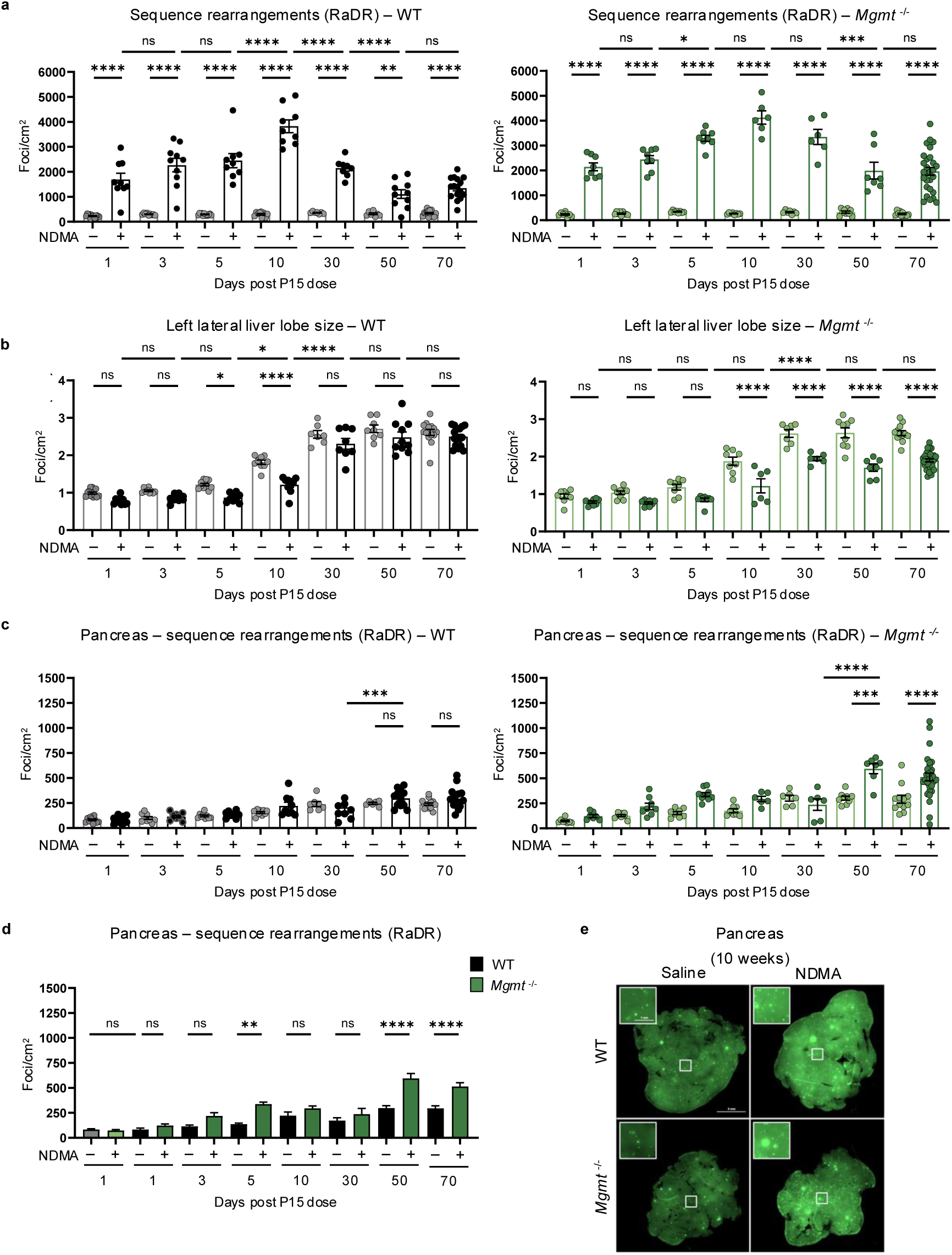
NDMA induces recombination events indicative of persistent genomic instability. **a**, Quantification of sequence rearrangement mutations in liver samples of WT (left, gray/black bars) and *Mgmt*^−/−^ (right, green bars) mice following NDMA or saline exposure at multiple timepoints, shows a peak at 10 days. RaDR foci (eGFP expression) per cm² quantified using machine learning image analysis. *n* ≥ 10 per group. **b**, Measurements of total left lateral liver lobe size (cm²) in WT (left) and *Mgmt*^−/−^ (right) mice across NDMA and saline groups. Significant reductions are observed in NDMA-treated WT mice at 5 and 10 days, and in *Mgmt*^−/−^ mice from 10 to 70 days post-exposure. *n* ≥ 10 per group. **c**, Quantitation of sequence rearrangement mutations in the pancreas, shown for WT (left) and *Mgmt*^−/−^ (right) mice. RaDR foci per cm² quantified by machine learning. Significant increases are detected in *Mgmt*^−/−^ mice after NDMA exposure at 50- and 70-days post-dose, but not in WT controls. *n* ≥ 10 per group. **d**, Direct comparison of pancreatic sequence rearrangement mutations post-NDMA exposure reveals significantly increased RaDR foci per cm² in *Mgmt*^−/−^ (green bars) compared to WT (gray/black bars) at 50 and 70 days. *n* ≥ 10 per group. **e**, Representative whole-mount fluorescence images of pancreatic tissue from WT and *Mgmt*^−/−^ RaDR-GFP mice 70 days after NDMA or saline treatment. Insets show higher magnification (scale = 1 mm) of boxed regions (scale = 5 mm at 2x) where areas are with GFP-positive foci indicating mutations. Statistical comparisons performed using one-way ANOVA with Šídák’s multiple comparisons test (**a**–**d**). Data are presented as mean ± s.e.m. Statistical significance: *p < 0.05, **p < 0.01, ***p < 0.001, ****p < 0.0001. NS, not significant.

**Extended Data Fig. 7:**
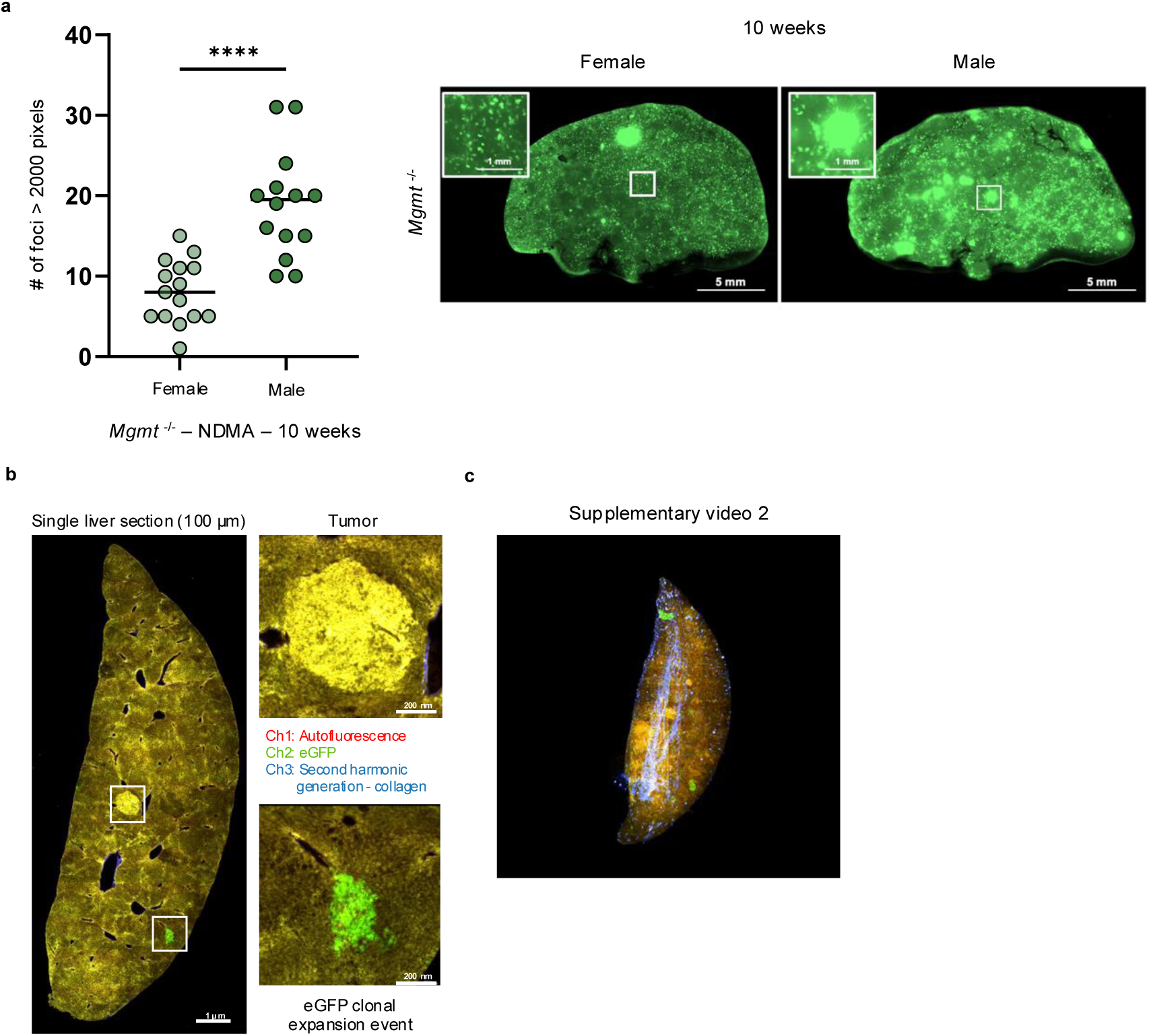
NDMA induces clonal expansion in *Mgmt*^−/−^ livers. **a**, At 10 weeks post-NDMA exposure large clonal expansion events are significantly greater in male *Mgmt*^−/−^ livers compared to females. A single large clonal expansion event was defined as RaDR-GFP foci exceeding 2000 pixels in area. Representative whole-mount eGFP images highlight the significant difference between female and male *Mgmt*^−/−^ livers. *n* ≥ 14 per group. Statistical comparisons performed using Mann-Whitney test. Data are presented as mean ± s.e.m. Statistical significance: *p < 0.05, **p < 0.01, ***p < 0.001, ****p < 0.0001. NS, not significant. **b**, 2-photon microscopy of a liver section (100 μm thick) from a ~13-month post-NDMA *Mgmt*^−/−^ female mouse visualizes both eGFP-positive clonal expansion events (green, lower boxes) and tumor areas detected by tissue autofluorescence (yellow, upper box). Insets display high-magnification images of the indicated regions. **c**, 3-dimensional reconstruction of sequential 2-photon images from (**b**) visualizes the spatial distribution of eGFP clonal expansion events (green) and tumors (yellow) within a *Mgmt*^−/−^ liver. The vascular network is displayed in blue. See Supplementary Video 2.

**Extended Data Fig. 8:**
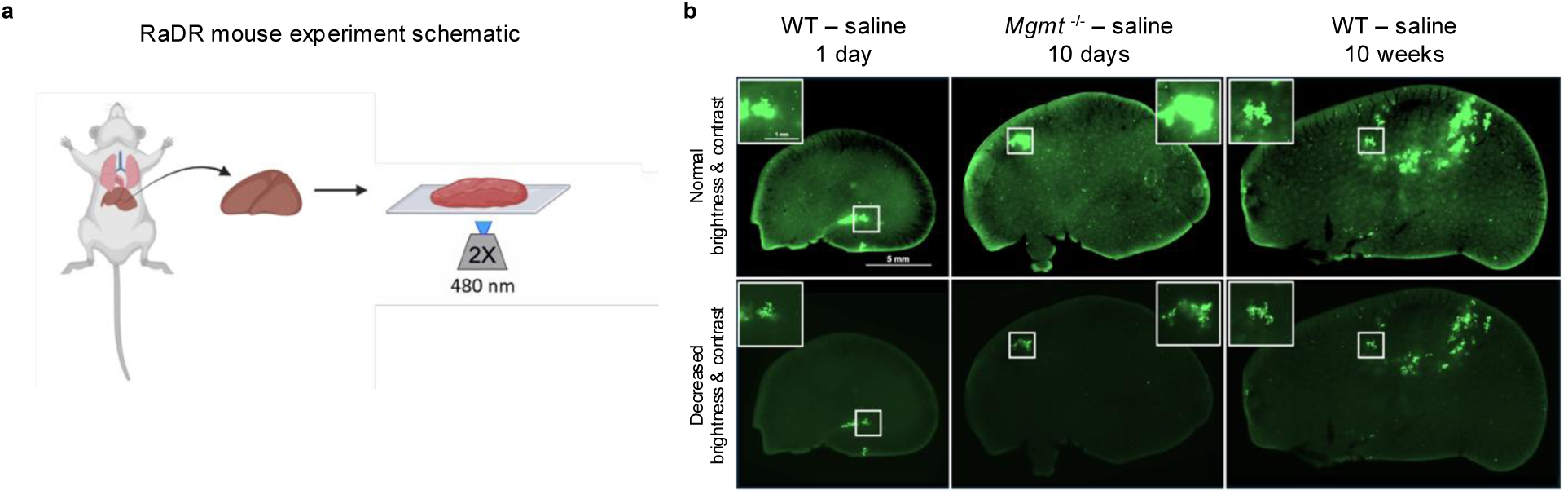
RaDR-GFP imaging. **a**, Schematic illustration of the RaDR-GFP mouse experiment. Freshly collected liver tissue is collected and mounted on a glass slide for GFP imaging at 2x magnification, with excitation at 480 nm. This protocol enables direct visualization of recombination reporter signals in liver tissue. **b**, Representative GFP fluorescence images of whole livers from WT and *Mgmt*^−/−^ RaDR mice at 1 day, 10 days, and 10 weeks post-saline treatment, imaged at consistent exposure and emission settings. In rare cases (~2%), spontaneous recombination of the direct repeat substrate occurs early in development which can lead to EGFP expression in many daughter cells. Early recombination events can be distinguished from the majority by the presence of a jagged edge with a decrease in brightness and contrast. Out of 327 mice, this was observed in 7 samples, and these were eliminated from analysis. Insets highlight regions of interest.

**Extended Data Fig. 9:**
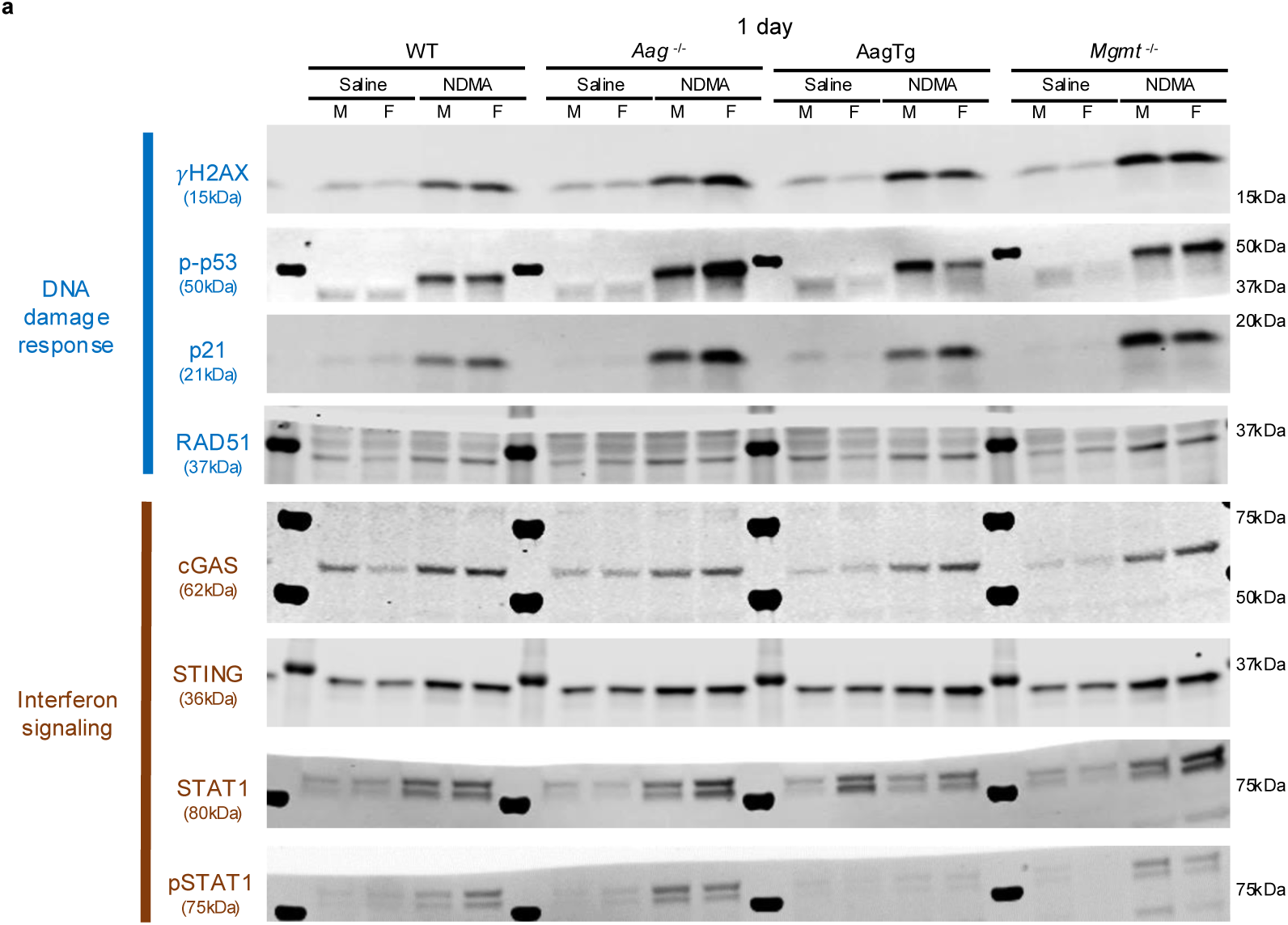
Western Blot Images.

**Extended Data Fig. 10:**
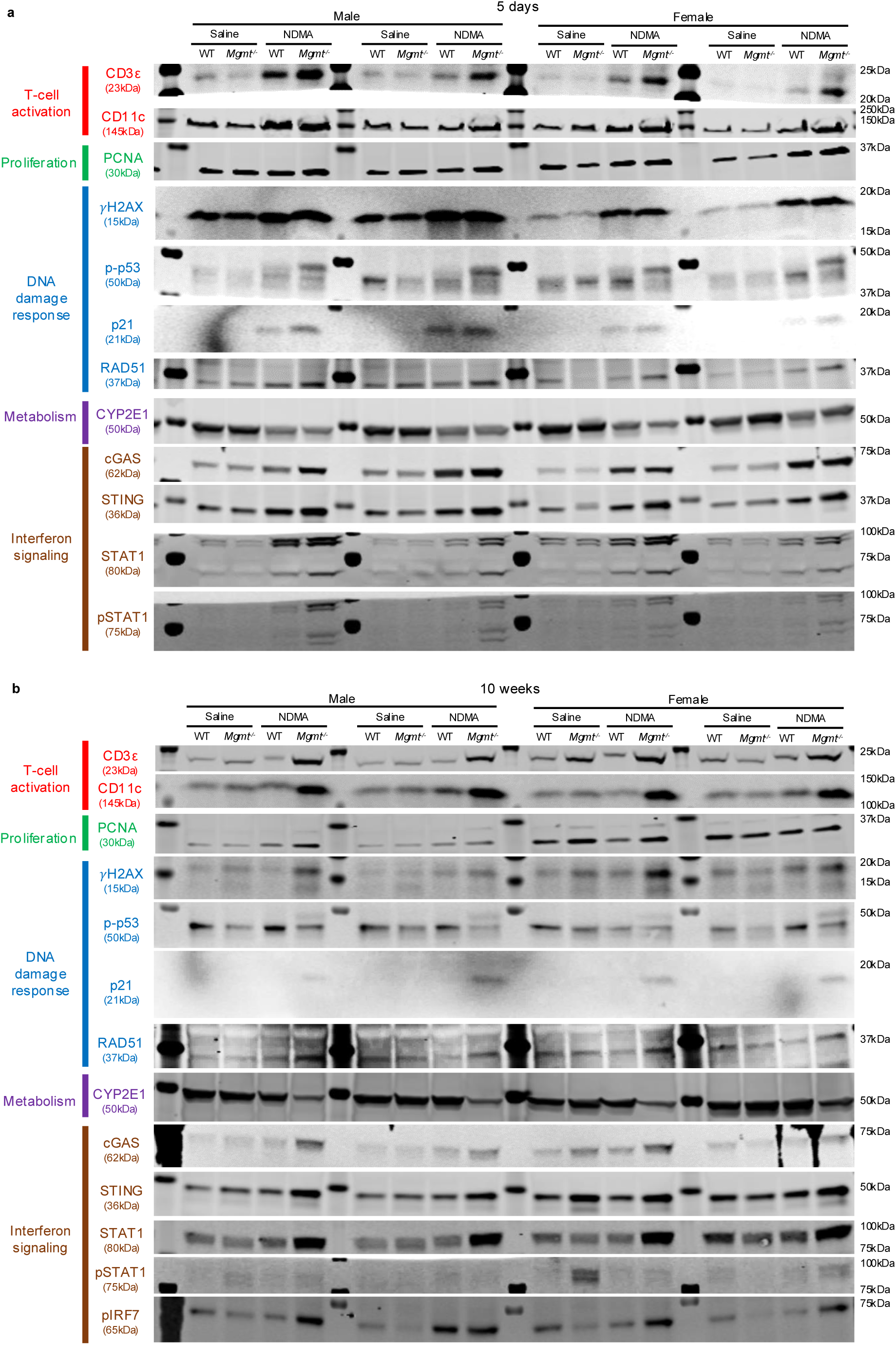
Western Blot Images.

## Notes

### Competing Interest Statement

The authors have declared no competing interest.

